# Patterns of and processes shaping population structure and introgression among recently differentiated *Drosophila melanogaster* populations

**DOI:** 10.1101/2021.06.25.449842

**Authors:** J.M. Coughlan, A.J. Dagilis, A. Serrato-Capuchina, H. Elias, D. Peede, K. Isbell, D.M. Castillo, B.S. Cooper, D.R. Matute

**Author notes:** Biology Department University of North Carolina, 250 Bell Tower Drive, Genome Sciences Building Chapel Hill, NC 27510. Denotes equal contribution (J.M. Coughlan, A.J. Dagilis). **Author contributions** **Experimental design:** JMC, AJD, DMC, and DRM designed all experiments and data analyses. **Data collection:** JMC, ASC, HE, DP, KI, BSC, and DRM collected the data, including field work, mate choice experiments, and DNA extractions. **Data analyses:** JMC and AJD performed all data analyses. **Manuscript writing:** JMC, AJD, DMC, BSC, and DRM wrote the manuscript.

## Abstract

Despite a century of genetic analysis, the evolutionary history underlying patterns of exceptional genetic and phenotypic variation in the model organism *Drosophila melanogaster* remains poorly understood. How genetic and phenotypic variation is partitioned across the range of *D. melanogaster*, particularly in its putative ancestral range in Subtropical Africa, remains unresolved. Here, we assess patterns of population genetic structure, admixture, mate preference, and genetic incompatibility across a global sample, including 174 new accessions from remote regions within Subtropical Africa. While almost all Out of Africa genomes correspond to a single genetic ancestry, different geographic regions within Africa contain multiple ancestries, with substantial cryptic diversity in Subtropical Africa. Admixture between distinct lineages is prevalent across the range, but admixture rates vary between lineages. Female mate choice within Subtropical Africa is highly polymorphic and behavioral types are not monophyletic. The genetic architecture of mate choice is highly polygenic, including loci associated with neurological development, behavior, olfactory perception, and learning. Finally, we discovered that many segregating putative incompatibilities likely evolved during or after expansion out of Africa. This work contributes to our understanding of the evolutionary history of a key model system, and provides insight into the distribution of polymorphic reproductive barriers.

## INTRODUCTION

*Drosophila melanogaster* has remained one of the most powerful genetic systems to understand the molecular underpinnings of animal biology since its development in the early 20th century [1–3]. The species is distributed globally and is commonly associated with human settlements [4, 5]. Recent sampling efforts have strongly suggested that *D. melanogaster* originated in the African mopane forest and initially bred on marula fruits, with a transition to human commensalism within Africa approximately 10,000-13,000 years ago [4, 5]. Yet, much remains unknown about the natural history, distribution, and evolutionary history of *D. melanogaster*, particularly in its ancestral range. Given the importance of model systems, like *D. melanogaster*, to our understanding of the genetic basis of morphological [6, 7], physiological [8], and behavioral [9] traits, as well as our understanding of different evolutionary processes in both natural and experimental contexts [10–12], it remains critical to understand how genetic and phenotypic variation evolved and is maintained in the ancestral range.

Although significant population genetic structure between Africa and non-African populations has long been recognized in *D. melanogaster* [13], and the existence of population genetic structure within Africa has more recently been suggested [14–16], the partitioning of genetic diversity within the ancestral range is largely still unresolved. Early multilocus surveys found limited to modest structure within Africa [14, 17], and supported distinct West and East African clades [15]. Most recent efforts suggest modern day remnants of ancestral *D. melanogaster* lineages exist as isolated, genetically unique populations within the putative ancestral range [5,16,18]. Human-aided migration following the transition to human commensalism is thought to have contributed to both the within Africa expansion [4,5,19], and subsequent global expansion, the latter of which likely resulted from a single out of African event [5,20–22], with multiple bottlenecks [23, 24]. However, despite housing the vast majority of genetic diversity, the demographic processes shaping population genetic structure in the ancestral range remain largely unknown (though see [4, 5]).

Post expansion, multiple historical events have also created opportunities for human-mediated admixture between genetically distinct lineages of *D. melanogaster*. It has previously been suggested that the opening of western commercial routes and the ‘Scramble for Africa’ facilitated hybridization between local African populations of *D. melanogaster* and invading non-African *D. melanogaster* individuals, particularly in more urban areas [25, 26]. Indeed, the extent of non-African ancestry in Africa is widely variable between populations [18], with some evidence for more pronounced signatures of admixture in urban populations, [25, 26]. Second, human migration associated with slavery roughly 400 years ago produced a secondary contact zone between African and non-African populations of *D. melanogaster* in the southeastern United States and the Caribbean [27–29]. However, it is unknown whether cryptic genetic lineages within Africa [16] have differentially contributed African ancestry outside of Africa [30]. Furthermore, the extent to which admixture between Out of Africa and African lineages contributes to the modern-day genetic composition of *D. melanogaster* within Africa is relatively unresolved (but see [18,25,26]). Lastly, patterns of admixture between African populations are almost entirely unexplored, despite their relevance for unraveling the evolutionary history of *D. melanogaster*.

One mechanism that could contribute to differentiation among *D. melanogaster* populations is behavioral variation between populations which results in assortative mating. Behavioral surveys of female mate choice within *D. melanogaster* from Subtropical Africa revealed a potential case of incipient speciation within *D. melanogaster* ([13,31–33]. At least two more cases of region-specific female mate preference have also been described [26, 34]). Cosmopolitan (denoted ‘M’) flies are globally distributed, including within Subtropical Africa, while a second lineage is largely restricted to Zambia and Zimbabwe (denoted ‘Z’). While Z females show significant preference for Z males, M females show no preference, and males of both lineages court both types of females at similar rates [31,33,35]. Early attempts to map the genetic basis of female preference found the trait to be highly polygenic, involving loci on all major chromosome arms, and significant non-additive effects [31, 32]. Despite the contributions of *D. melanogaster* to the field of speciation [36–41], it remains unknown whether the frequencies of these behavioral types covary with patterns of *D. melanogaster* population structure within Subtropical Africa. Understanding if/how behavioral isolation is associated with genetic structuring can allow us to determine if divergence in complex, quantitative traits—such as mate choice—can contribute to stable genetic divergence within a species; a hypothesis which is still widely debated [42–44].

In addition to variation in female mate choice, negative epistatic interactions between polymorphic loci, similar to hybrid incompatibilities, can cause substantial variation in fitness, including the production of low fitness individuals within a species [45–49]. While it has been hypothesized that these incompatibilities are geographically structured across the *D. melanogaster* range [45], very little is known about the geographic origins or distributions of these alleles. In particular, it remains unknown if these alleles correspond to previously described behavioral and/or genetic lineages (i.e. [13,16,33]). Determining how these forms of reproductive isolation correspond with population structure and patterns of gene flow can contribute to our understanding of the early stages of population differentiation and speciation.

Here we address the extent of genetic differentiation and admixture in *D. melanogaster*, relate patterns of genetic structure to behavioral differences in the putative ancestral range, and describe the geographic distribution of putative incompatibility alleles and their contributions to admixture and population structure. This work leverages genome-wide information from 420 individuals, including 174 samples from novel and previously under-sampled geographic regions as well as extensive behavioral assays of flies from Subtropical Africa. Specifically, we ask three questions: (*i*) does *D. melanogaster* show population genetic structure in Africa, particularly within the ancestral range? (*i*) How much gene flow occurs between genetic lineages? And (*i*) how do two forms of polymorphic RI (mate choice and genetic incompatibilities) correlate with population structure in this system? Our results help to clarify the demographic history of *D. melanogaster* and provide some insights into the persistence of genetically unique clades within *D. melanogaster*.

## RESULTS

### Diversity, divergence, and evolutionary relationships among populations of *D. melanogaster*

To understand the global distribution of diversity and population structure of *D. melanogaster,* we combined whole genome resequence data from 420 lines of *D. melanogaster* from around the world, including 174 newly sequenced genomes from un- and under-sampled, rural locales within the proposed ancestral range in Subtropical Africa. PCA revealed that genetic variation within *D. melanogaster* is mainly structured between flies from Subtropical Africa and Out of Africa (OOA), as reflected by PC1 (which explains 46.7% of the variation). While somewhat intermediate along PC1, Ethiopia, as well as East and West Africa, are much more distinct from OOA and Subtropical Africa along PC2 (which explains 18.6% of the total variation). We further identified eight unique genetic ancestries using *K*-means clustering in *PCAngsd* [50]. Four of these ancestries predominantly occur in Subtropical Africa, three largely correspond to one each of Ethiopia, West Africa, and East Africa, and a final ancestry type is most common in all OOA accessions (including *D. melanogaster* from North Africa; Figure 1).

**FIGURE 1.**
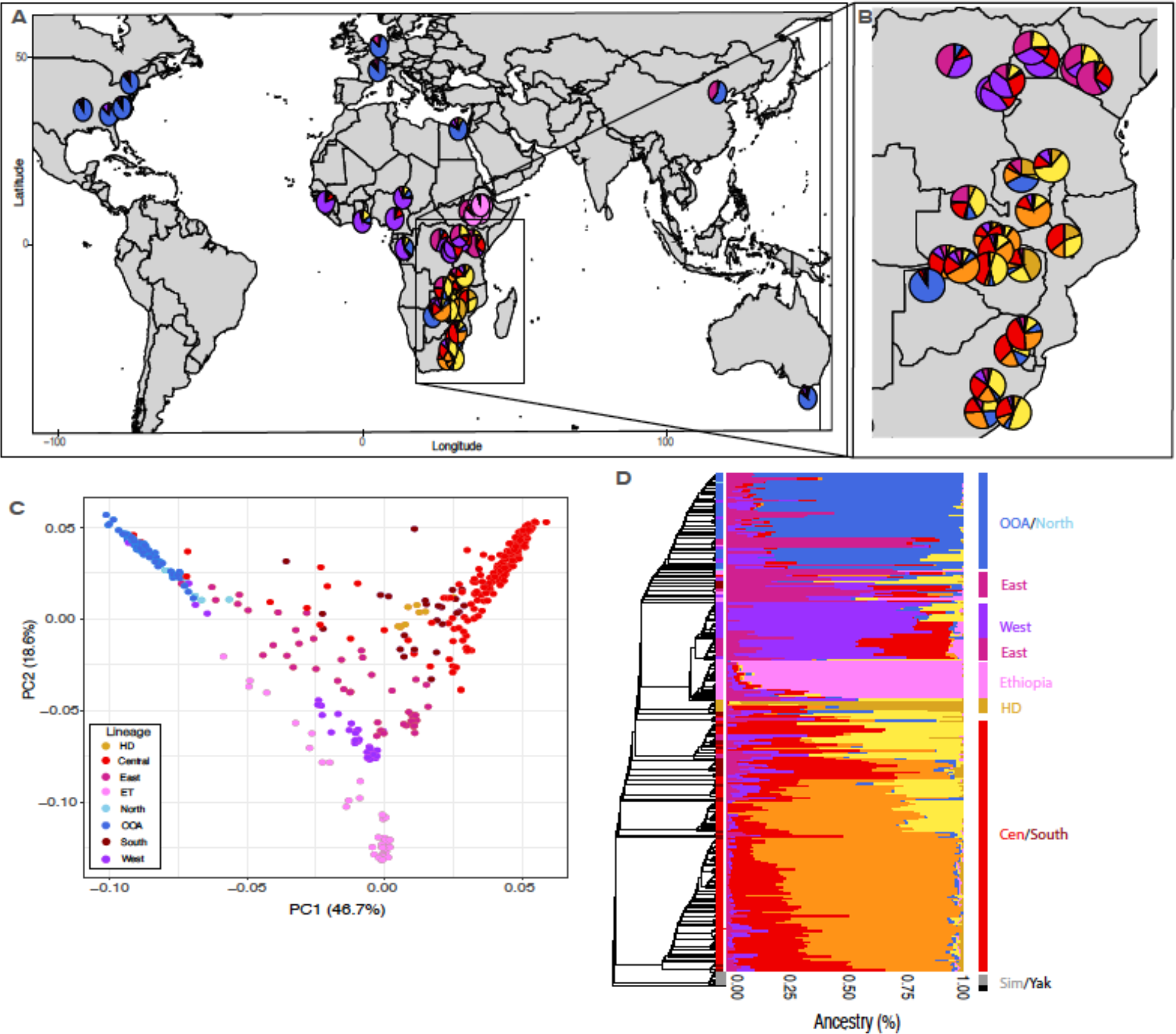
Broad sampling of Drosophila melanogaster from Subtropical Africa reveals substantial cryptic genetic structure. (**A**) Geographic sampling of 420 genomes from a global distribution of D. melanogaster; pie charts represent the average ancestry determined by PCAngsd from that sampling local. (**B**) Zoom-in panel from (A), focusing on Subtropical and East African accessions. (**C**) PCA of all genomes, colors indicate the genetic lineage of the sample, defined herein. We also denote samples from North and South Africa (although they reside within OOA and Southern Africa, respectively). (**D**) Phylogeny of all accessions with an ancestry plot denoting their average genomic composition of each sample. Labels to the right of the ancestry plot indicate how samples cluster into the six genetic lineages that we identify using PCA, K- means clustering, and phylogenetic reconstruction.

*K*-means clustering and PC analyses largely agree with our consensus phylogeny with *D. simulans* and *D. yakuba* as outgroups. Of the eight ancestry types identified, four correspond to largely monophyletic clades; OOA, Ethiopia, West Africa, and a unique ancestry type from Subtropical Africa that comprises nine individuals predominantly from Harare, Zimbabwe (denoted by gold in Figure 1). We refer to this lineage as Harare-Distinct (HD). Of the remaining four ancestries, individuals from East Africa cluster together in a PCA, but are split across our phylogeny, with some individuals sister to West Africa, and others sister to all OOA individuals. The remaining three ancestries are largely found in individuals from Subtropical Africa, but do not correspond to monophyletic clades or unique sampling locales. They also do not exist in pure form; the vast majority of Subtropical African individuals comprise two, and sometimes three of these ancestries (denoted as yellow, red, and orange in Figure 1). Although ancestry types do not correspond to monophyletic groups in these Subtropical African individuals, we note that the largest and most distantly related clade of Subtropical African flies has the largest proportion of one ancestry type (indicated by orange in Figure 1), and largely contain samples from more remote sampling locales. Combining these results, we define six genetic lineages that we use for subsequent analyses: OOA, Ethiopia, East Africa, West Africa, HD, and all other Subtropical African flies (which we refer to as Southern Africa).

Despite substantial population genetic structure, global *F_ST_* estimates were relatively low in almost every comparison, with the lowest global *F*_ST_ found between East and West Africa (*F*_ST_=0.027), and the highest between HD and OOA (*F_ST_*=0.241; Table S2). Variation in pairwise global *F_ST_* is likely due to differences in nucleotide diversity, as all pairwise comparisons of *D*_xy_ were very similar (Table S2), and the populations with the highest global *F*_ST_ were those in which both populations also showed the lowest pairwise nucleotide diversity (*π*; Table S2; Figure 2).

**FIGURE 2.**
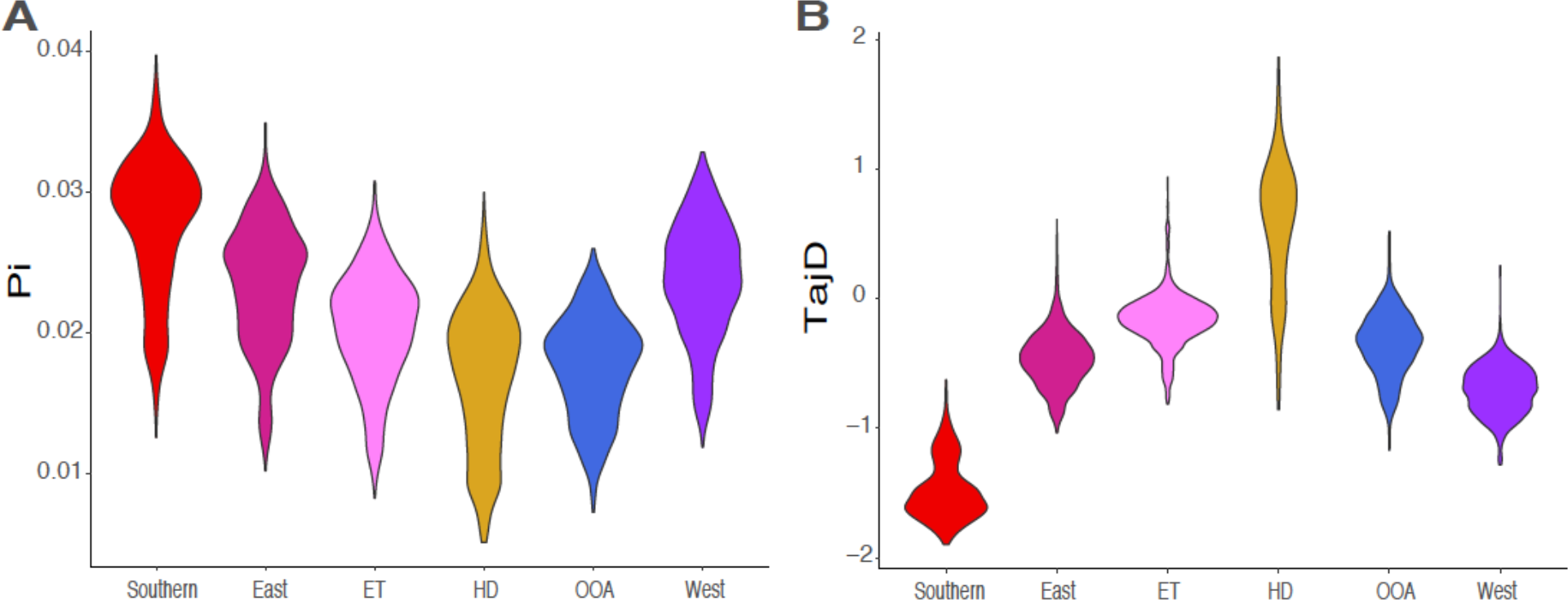
Genome-wide statistics for each genetic lineage. Average (**A**) nucleotide diversity (π) and (**B**) Tajima’s D for each genetic lineage.

In addition to substantial cryptic genetic structure, the Southern African lineage contains substantially higher levels of genetic diversity than any other genetic lineage (Figure 2). This diversity is likely caused by an excess of rare alleles, as the Southern African lineage also shows the most negative values of Tajima’s D, and a left-skewed Site-Frequency Spectrum (SFS), in line with this lineage having a much larger effective population size and a recent history of population expansion (Figure 2, Figure S9; as suggested in [22]). In contrast, two lineages show signals of recent population contraction and lower levels of diversity: OOA (including North Africa) and HD (Figure 2). Tajima’s D and the SFS also differ between chromosomes, although the direction of these differences depended on the genetic lineage (based on an ANOVA with Type III SS: chromosome arm × genetic lineage effect: *F*=162.9, *df*=20, *p*<0.0001). For four lineages, the *X* chromosome had lower values of Tajima’s D (Southern Africa, East Africa, West Africa, and HD), while Ethiopia and OOA show the opposite pattern (Figure 2; Figure S3; Figure S9). Overall, these results suggest that both different lineages, and different chromosome arms, of *D. melanogaster* have experienced different demographic histories and even within Subtropical Africa, the HD lineage has a significantly different demographic history than the rest of Southern Africa.

### Patterns of gene flow throughout the range of *D. melanogaster*

We next evaluated the extent of gene flow among distinct lineages within a global sampling of *D. melanogaster*. Specifically, we focus on three potential cases to better understand the sources and dynamics of gene flow across the range of *D. melanogaster*: (1) between Subtropical Africa and other African lineages, (2) the extent of gene flow between OOA and both HD and Southern Africa, and (3) the source(s) of African ancestry in the SE United States (as proposed by [27, 28]). We describe the results for each of these cases as follows.

First, we evaluated the extent of admixture among genetic lineages within Africa. We find evidence for extensive gene exchange between Southern Africa and both East and West Africa, but not between Southern Africa and either Ethiopia or HD (Table S3, Figure 3). Although gene flow between Southern and both East and West Africa is pervasive, the magnitude of gene flow is likely quite low, as only 1-3% of the genome is inferred to be admixed between Southern Africa and each of East and West Africa, respectively (depending on whether *f*_G_ or *f*_dM_ is used; Table S3, Figure 3). Similarly, we find significant gene flow between HD and both East and West Africa (Table S3, Figure 3). Despite closer geographic proximity between most East African samples and Southern African samples, introgression between West Africa and both Southern Africa and HD is significantly elevated compared to introgression between East Africa and either Southern Africa or HD (Figure 3; *f*_G_ between West Africa and either HD or Southern Africa ranged from 0.02-0.033, while *f*_G_ between East Africa and either HD or Southern Africa ranged from 0.017-0.022; we find no significant difference between Southern Africa and HD in the extent of gene flow with either West or East Africa). Overall, this suggests that the magnitude of admixture is not equivalent between Subtropical African and other African *D. melanogaster* lineages and may not simply correspond to geographic distance.

**FIGURE 3:**
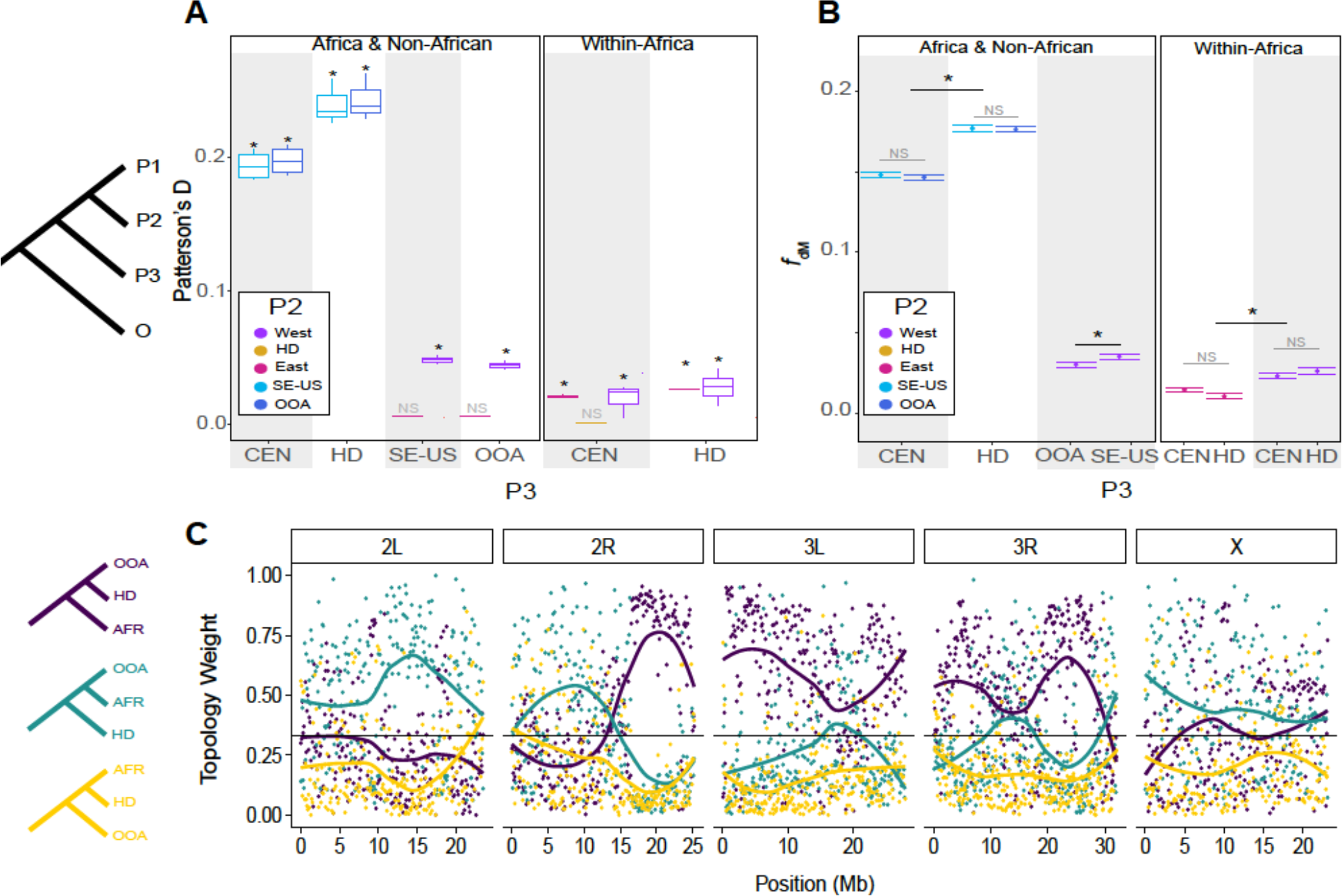
Patterns of gene flow between different African and non-African lineages of D. melanogaster. (**A**) Boxplots of Patterson’s D for all trios given the phylogeny (((OOA, ((East, West),Ethiopia),HD), Southern Africa). P3 is given on the X- axis, with the identity of the P2 population denoted by the color of the boxplot. Values of D represent the range of values over multiple P1 populations. Comparisons that yielded a significant Patterson’s D based on a standard block jackknife procedure [54] are denoted with an asterisk, otherwise they were deemed Non-Significant (NS). (**B**) Mean and standard errors of 20-SNP windows of f_dM_ from across the genome for each African genetic lineage showing significant introgression with either other African or non-Africa lineages. The OOA lineage is split into those from the SE United States (SE-US) and all others (OOA). Significance was determined by *ANOVA*s with Type III SS: NS= Not Significant, * = 0.01<p<0.05. (**C**) Weighted topologies for three configurations of OOA, HD, and Africa (denoted by colored phylogenies to the left).

Second, we evaluated the extent of introgression between OOA and both Southern Africa and HD. Specifically, we aimed to test whether back migration from OOA may be responsible for the genetically unique HD lineage (denoted in gold in Figures 1 and 2) as a test of whether urbanization has facilitated introgression in Africa. Other genomic evidence suggests that this may be the case- the HD lineage shows substantially reduced diversity and genomic patterns of a bottleneck relative to Southern Africa, despite no geographic separation (Figure 1, Figure 2), and mimics patterns of diversity and Tajima’s D seen in OOA populations (Figure 2). In line with this hypothesis, we find evidence of admixture between HD and OOA; *f*_dM_ is elevated between OOA and HD when compared to OOA and Southern Africa (Figure 3), wherein 18-20% of OOA and HD genomes have introgressed, while only 15% of genomes between OOA and Southern Africa have introgressed (depending on whether *f*_G_ or *f*_dM_ is used; Table S3, Figure 3). However, tree topology weights generated by *twisst* further reveal that the unique HD lineage does not appear to be simply a hybrid of flies from Southern Africa and OOA (Figure 3C). While large proportions of the genome show near complete support for topologies in which HD is sister to OOA (purple, in Figure 3C), the second most common topology does not place HD sister to samples from Africa, but rather places all other African samples sister to OOA (teal, Figure 3C). This suggests that rather than being a patchwork of African and OOA ancestry (in which we expect yellow and purple topologies to be common in Figure 3C), HD carries distinct genetic variation, at least in genomic regions where the topology supports all other African genomes sister to OOA (i.e. the center of 2L). Extensive introgression between African lineages and OOA may also contribute to these patterns. In total, this suggests that while HD has likely experienced substantial introgression with OOA, it has also likely experienced a unique evolutionary history from either OOA or the rest of Southern Africa.

Last, we studied the origin of the African alleles harbored in the SE United States. We find significantly elevated Patterson’s *D* between SE United States and each of Southern Africa, West Africa, and HD which suggests these populations have contributed to *D. melanogaster* from the SE United States. We do not observe a similar pattern for Ethiopia or East Africa (Table S3, Figure 3). Patterns of introgression are not unique to the SE United States, as we find elevated Patterson’s *D* between all other OOA lines and each of Southern Africa, West Africa, and HD (Table S3, Figure 3), suggesting that the contribution of these African populations might precede the split of different OOA populations. However, if *D. melanogaster* from the SE United States experienced a second pulse of introgression unique from other OOA populations, then we predict that SE United States flies should show elevated signals of introgression with at least one African source relative to other OOA flies. In line with this prediction, both *f*_G_ and *f*_dM_ between West Africa and the SE United States are elevated relative to either *f*_G_ and *f*_dM_ between West Africa and other OOA populations (Table S3, Figure 3), while Southern Africa and HD show no difference in either *f*_G_ and *f*_dM_ with the SE United States or other other OOA lines (Table S3, Figure 3). However, it is challenging to detect regions of the genome that show unique signals of introgression between West Africa and the SE United States relative to West Africa and all other OOA accessions, as the landscape of *f*_dM_ across the genome is highly correlated between these two comparisons (*r^2^*=0.59, *p*<0.001; as it is between Southern Africa and OOA and Southern Africa and SE United States: *r^2^*=0.509, *p*<0.001, and between HD and OOA and HD and SE United States: *r^2^*=0.607, *p*<0.0001). Moreover, the weighted topologies with West Africa as sister to all OOA versus only accessions from the SE United States are highly similar (Figure S7). The lack of distinction may also be caused by rapid purging of introgressed alleles, reducing any signals of introgression in a short time- span [51]. In total, this suggests that all OOA populations have experienced some level of gene flow with Southern Africa, West Africa, and HD, but flies from the SE United States may have experienced a second, and independent pulse of West African ancestry (as has been hypothesized by [27, 28]).

We next examined how patterns of introgression vary across the genome within Africa and between African and non-African lineages. For these analyses, we focus on the three scenarios outlined above: (1) Introgression within Africa (as measured between West Africa and Southern Africa), (2) introgression between OOA and the unique HD lineage, and (3) introgression between West Africa and the SE United States. In all instances, Ethiopia was used as P1 as we find little to no evidence of admixture between Ethiopia and any P3 used herein (Table S3). We find that chromosomes significantly differ in the extent of introgression for all comparisons (as measured by *f*_dM_; West-SE-US: *F*=47.11, *df*=5, *p*<0.001; West-South: *F*=18.36, *df*=5, *p*<0.001; OOA-HD: *F*=16.01, *df*=5, *p*<0.001). For all three comparisons, the *X* chromosome showed significantly higher *f*_dM_ values, than almost every autosome (Table S4). However, definitive evidence of increased introgression on the *X-*chromosome is much less apparent when using weighted topologies (Figure 3, Figure S7).

Many genes which may be involved in the transition to human commensalism fall within the top 1% of *f*_dM_ windows (see Table S5 for full list). For example, these windows include several genes involved in insecticide resistance, (including *Cyp6a18* (between OOA-HD), *Cyp313a* and *ACE* (West-SE-US), and *Cyp12a4* and *LRR* (West-South)), several genes related to metabolism, feeding behavior, and perception of and response to food sources (including *happyhour*, *pbx, for,* and *Gfat1* (West-South), *NPFR, lovit,* and *Ald1* (West-SE-US), and *Gr5a* (OOA-HD)), as well as genes involved in immune function (*Dcr-2*, *Rab4*, *DptA*, *DptB*, and *Tl* (OOA-HD), *Npc2h*, *Npc2g*, and *ben* (West-South), and *Charon* and *Ance-2* (West-Se-US)). Although causative connections are still needed, it is perhaps unsurprising that alleles putatively related to human commensalism may be overrepresented in three examples of potentially long-range, and potentially human-mediated admixture.

Lastly, we find several genes associated with mating behavior fall within the top 1% of introgressed regions including *Desat2* (West-SE-US; although we note that this gene may also be involved in desiccation resistance, as is the case with other cuticular hydrocarbons [52, 53]), *fru* (West-South), and *5-HT1A*, *clt*, and *Oamb* (OOA-HD). While these regions may not be involved in human commensalism *per se*, introgression at these loci may have implications for the distribution of female mate choice, and the distribution of female mate choice may be influenced by human commensalism; a possibility that we explore below.

### Patterns of female mate choice in Subtropical Africa

To understand if and how strong female mate preference originally described in Subtropical Africa corresponds to any of the Subtropical African ancestries that we identify herein, we surveyed a subset of lines from Subtropical Africa for the existence of female mate choice using replicated choice experiments. Briefly, 47 focal isofemales from ten sampling locales in Subtropical Africa were presented with both a Subtropical Africa Z (ZS2) and OOA M (RAL371) isoline male. We quantified female mate preference as a significant deviation from random choice, wherein ZS2 males were more likely to copulate than RAL371 males. We find that female mate preference was common in our experiment, but far from fixed across Subtropical Africa (e.g. 19/47 lines had a preference for ZS2 males; Table S6, Figure S4). The frequency of lines showing female mate preference varied among sampling locales, with the average proportion of females exhibiting preference per population being 44% (range: 0-100%; *X^2^*=19.04, *df*=9, *p*=0.025; Figure 4; Table S6). Populations also varied in the strength of preference. Some populations had nearly complete preference for Z males, and others showed a mild aversion to Z males (*X^2^*=18.63, *df*=9, *p*=0.029; Table S6; Figure S4; average proportion of Z males chosen per population ranged from 0.398-0.969). Lastly, we find that isolines that exhibit female mate choice are not monophyletic, nor do any major PCs ascribing genetic variation among phenotyped flies separate isolines with female preference from those that do not (Figure S5). Our results suggest that the dynamics of mate preference are complex with Subtropical Africa; both the presence and strength of female mate choice vary within and among populations, and the presence of mate preference does not correspond with any obvious genetic structure.

**FIGURE 4:**
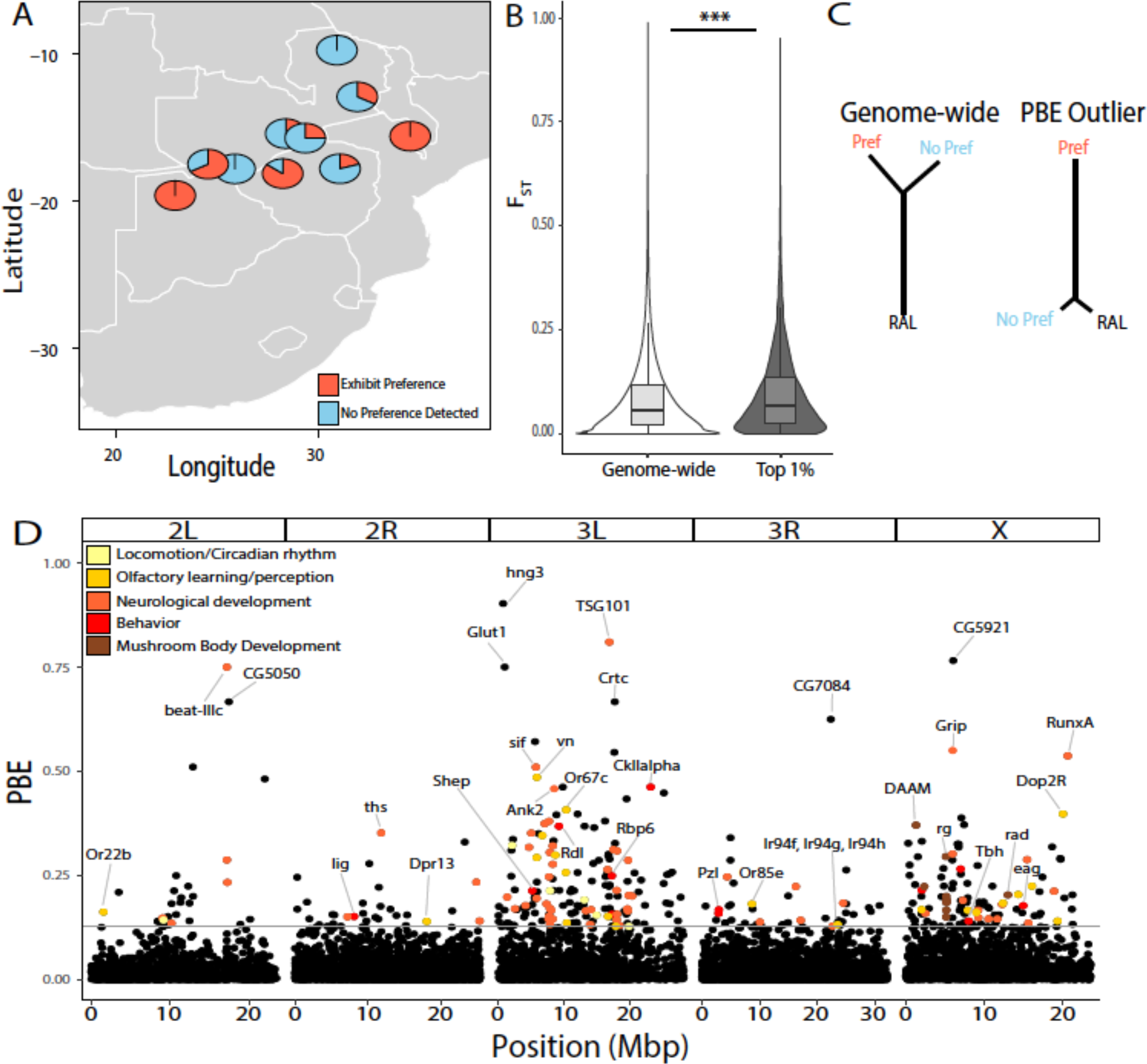
Geographic and genetic dissection of female preference in Subtropical Africa. (**A**) Frequency of isofemale lines that show significant female mate choice in two-male choice experiments per sampling locale. (**B**) Average F_ST_ is elevated in the top 1% PBE regions between the Southern Africa lineage (regardless of mate preference phenotype) and all other geographic regions. (**C**) Cartoon illustrating that the PBE statistic can infer branch-specific evolution of the focal population (in this case, flies with significant female mate preference (red, “pref”), relative to flies with no preference (blue, “no pref”) and a third population of RAL)). (**D**) PBE of D. melanogaster lines with strong mate preference, measured in 1KB windows across the five major chromosome arms of D. melanogaster. Grey horizontal line indicates the top 1% threshold. Colored dots indicate a sample of genes with functions relevant to female mate preference.

We leveraged the natural diversity in female mate choice among these 47 lines from Subtropical Africa to determine what regions of the genome are associated with female mate choice. Given that isolines with strong preference do not show significant population structure relative to isolines with no strong preference (Figure S5), we instead quantified branch-specific evolution for lines with significant female mate preference. We calculated the population branch excess (*PBE*) statistic [55], with Subtropical African lines with strong female mate preference as the focal population and Subtropical African lines with no strong female mate choice and RAL as the non-focal populations. We defined the top 1% (368 windows) as outliers. These regions represent all 5 chromosomal arms, although there is a significant excess of outliers on 3L and on the *X* chromosome (*X^2^*=183.01, df=4, p<0.0001). Of these 368 windows, 283 contain at least one gene, and 84 of the remaining 85 windows are proximal to at least one gene (of the windows in which genes are proximal, genes occur within 16.2KB on average, range: 2.5-100KB). A total of 407 unique genes are included either within these windows, or in close proximity (i.e. within 16.2KB on average). Gene Ontology overrepresentation analyses indicate that of these 407 genes, there is an excess of genes associated with learning, memory, cognition, chemotaxis, sensory perception, detection of stimulus, behavior, and several categories associated with neurological development (as well as other biological functions; see Table S8 for details).

Several individual genes are of potential interest for future study (Table S7). We find thirty genes that are known to influence behavior in the top 1% of *PBE* outliers, including five genes specifically involved in courtship behavior (*Shep, Rdl*, *Tbh, lig*, and *eag*). Thirty-eight genes that are involved in sensory perception and sensory learning are also within the top 1% of *PBE* values, particularly those involved in sensory perception of smell or olfactory learning, including three odorant receptor genes (*Or22b*, *Or67c*, and *Or85e*), four defective proboscis extension response genes (*dpr8*, *dpr13*, *dpr14*, *dpr18*), seven ionotropic receptor genes (*Ir11a*, *Ir67b*, *Ir67c*, *Ir75a*, *Ir75b*, *Ir94f*, *Ir94h*), *Dop2R* and *vn* (both involved in olfactory learning), as well as *Ank2* and *sif,* which are involved in sensory perception of sound and visual perception, respectively. Five genes which are known to influence male aggression fall in the top 1% of behavioral *PBE* values (*Dop2R, Rpb6, CkIIalpha, Tbh,* and *Rdl*). Z males tend to be substantially more aggressive during courtship than their M male counterparts, particularly to Z females (Figure S8). Lastly, ten genes that are involved in mushroom body development are also included in the top 1% of *PBE* values, including *Frl, DAAM, PsGEF*, *rg*, and *rad*. Mushroom bodies play a central role in learning and memory, particularly of olfactory perception, and therefore may be important in differentiating male mates. Furthermore, mutants with aberrant mushroom bodies and inhibition of mushroom bodies can cause virgin females to reject matings [56, 57]. In total, we find that *D. melanogaster* from Subtropical Africa with significant mate preference differ in many regions across the genome, indicative of a polygenic basis of female mate choice.

We next sought to assess whether loci putatively involved in female mate choice were also more highly differentiated among a broader geographic sampling of *D. melanogaster* as a test of polygenic adaptation of this trait. Specifically, we asked whether behavioral *PBE* outliers were among the most highly differentiated loci between all Southern Africa lines (regardless of phenotype) and other geographic regions. We find that *F_ST_* among behavior outliers is elevated between Southern Africa and all other genetic lineages relative to genome-wide *F_ST_* (locus type (e.g. genome-wide versus *PBE* outlier): F=6.426, df=1, p=0.011; genetic lineage effect: F=23799.7, df=4, p<0.0001, locus type × lineage interaction: F=0.94, df=4, p=0.43, Figure 4, Figure S6). However, despite this modest evidence for selection, we do not find that *PBE* outlier loci are less likely to introgress than the rest of the genome, as would be expected if female mate choice served as a significant barrier to introgression (*f_dM_* for behavioral *PBE* outliers is not significantly reduced relative to the rest of the genome between Southern Africa and either West Africa: *t*=-0.064, *df*=406.27, *p*=0.95 or Europe: *t*=-1.52, *df*=428.69, *p*=0.13). Furthermore, individual isolines that differ in the strength of female mate preference show no difference in the history of introgression experienced (mean *f_G_* for lines with female mate choice = 0.097, mean *f_G_* for lines with no strong female mate choice= 0.112; *t*=-0.27, *df*=9.7, *p*=0.79). In total, this work suggests that while female mate choice may show modest signals of polygenic adaptation, it is likely not a very effective barrier to gene flow in natural *D. melanogaster* populations.

### Distribution of putative incompatibilities throughout D. melanogaster

We last assessed the distribution of alleles potentially involved in negative epistatic fitness effects in admixed populations (herein ‘incompatibility loci’) throughout the range of *D. melanogaster.* Although we refer to these loci as incompatibilities, they include any loci that are found in repulsion of one another, and thus may include traditional incompatibilities (i.e., involved in intrinsic postzygotic isolation), alleles involved in ecological hybrid breakdown, as well as loci involved in assortative mating [58]. We find that loci identified by [45], show slightly increased overall levels of differentiation than the genome-wide average between Southern and West Africa, but significantly lower differentiation than the genome-wide average between Europe and each of Southern and West Africa (Figure 5). In contrast, we find no significant difference in *F*_ST_ between loci identified by [46] and the genome-wide average for any comparison. Thus, while on average, putatively incompatible loci are slightly more differentiated between West and Southern Africa than other loci, they are not broadly differentiated on a global scale.

**FIGURE 5:**
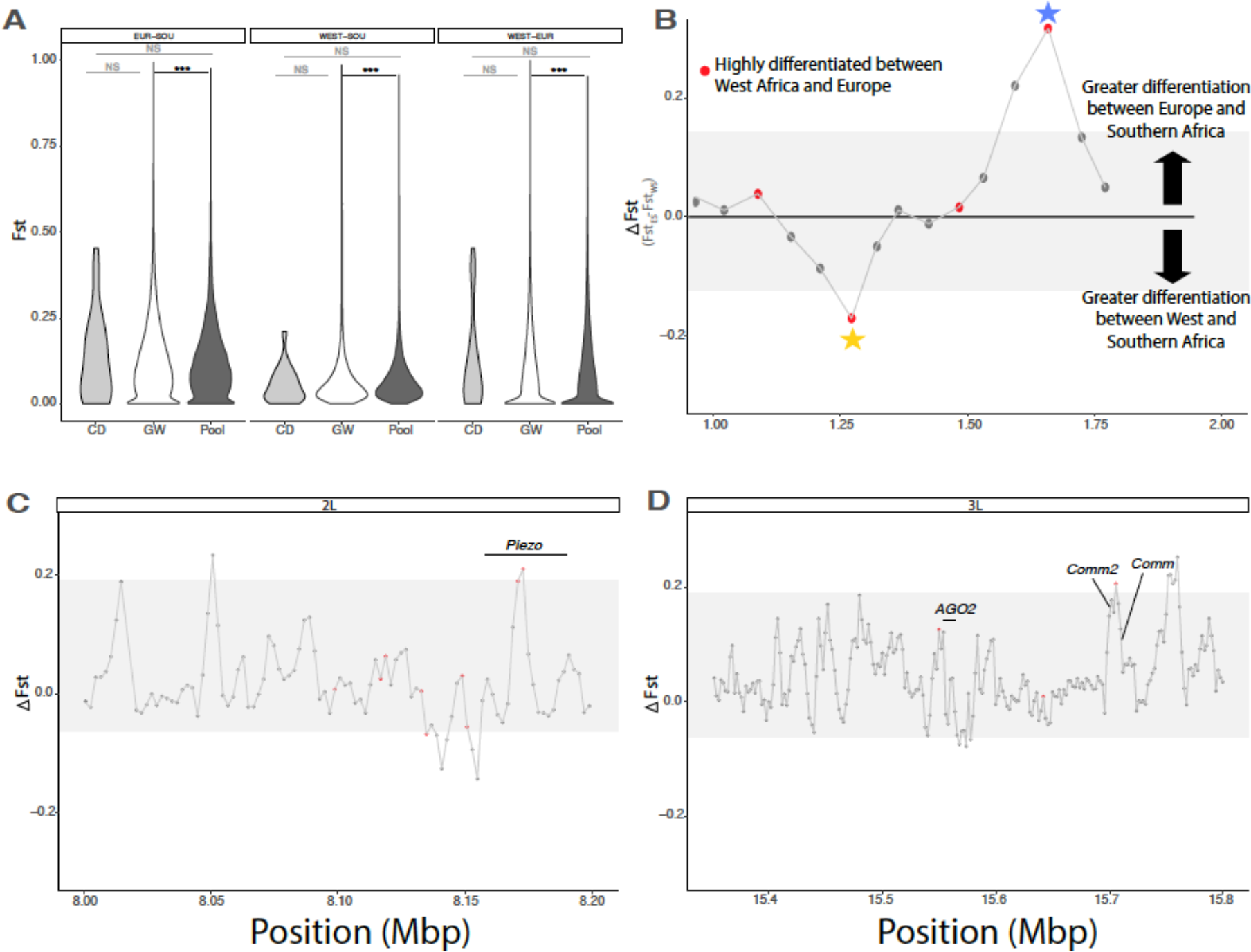
Global differentiation of putative incompatibility loci. (**A**) Distribution of F_ST_ for incompatibility loci from CD [46], Pool [45] and genome-wide for three comparisons: Southern Africa vs Europe, West Africa vs Southern Africa, and West Africa vs Europe. (**B**) Cartoon depiction of a scan for incompatibility loci that segregate between African lineages (yellow star) or likely arose during or after the Out of Africa expansion (blue star). The Y-axis denotes the difference in F_ST_ between Europe and Southern African (ES) and West and Southern Africa (WS), with more positive values indicating that F_ST_ is greater between Europe and Southern Africa than within Africa. Red points denote loci that are also highly differentiated between Europe and West Africa. Loci that are highly differentiated between Europe and both African lineages are putatively newer incompatibilities, while loci that are highly differentiated within Africa, but shared between Southern Africa and Europe may represent older incompatibilities. (**C,D**) Two zoomed in windows representing one incompatibility pair, as identified by Pool [45]. In panel (C) differentiation is elevated between Europe and Southern Africa and West Africa and Europe (but not West and Southern Africa) for loci within the gene *Piezo*, indicating this allele may be more recently derived in OOA populations. In (D) we show the corresponding locus: again, differentiation is elevated between Europe and each of West and Southern Africa (but not within Africa) in a window containing no genes, but slight up and down stream of *comm* and *comm2*, respectively. We also highlight the approximate position of *AGO2*, the hypothesized interacting partner (based on [45]), and show there is no increased divergence in this region.

We next assessed whether specific pairs of incompatibility loci showed elevated differentiation, and used patterns of differentiation to characterize their potential geographic origins. Out of 445 putative incompatibility loci identified by [45] and 45 putative incompatibility loci identified by [46], we identified only eight pairs of interacting incompatibility alleles with high differentiation within Africa, and low differentiation between Southern Africa and Europe; indicative of potentially older incompatibilities that divide genetic lineages within Africa. Within these eight loci, eight unique genes are included in the *F*_ST_ outlier windows, two of which were also found in our *PBE* behavioral analysis (*Abl,* and *eag;* Table S9). In contrast, we identified 71 pairs of loci with a signature of more recent origin (i.e. low differentiation in Africa, high differentiation between Europe and both West and Southern Africa). We find that the *F*_ST_ peaks within these 71 pairs of loci contain 133 unique genes (Table S9), 13 of which are also behavioral *PBE* outliers. These include *DAAM, Rbp6, beat-IIIC, Doa, Lasp, luna, Eip75, Mp, olf413, Pex7, pk, Svil,* and *eag* (we note this is a separate 1KB window than the window which shows a signal of an putatively older incompatibility, described immediately above). Many of these genes are involved in neurological development and/or behavior (*DAAM, Rbp6, eag, beat-IIIC, Eip75, pk,* and *Mb*), and reproduction (*Doa, Lasp*). Thus, many putative incompatibilities show a signature of more recent evolution, and a significant proportion of these loci may also be differentiated between behavioral types in Subtropical Africa.

## DISCUSSION

The evolutionary history of genetic model systems has been the target of extensive research, including *D. melanogaster* [4,5,16,18]. Nonetheless, sampling gaps across critical regions of its range have left crucial aspects of its history unexplored. We explore the partitioning of genetic diversity and patterns of gene flow across the global range of *D. melanogaster,* including 174 samples from previously under-sampled regions within the ancestral range. Furthermore, we evaluate the distribution of two sources of polymorphic reproductive isolation that have been previously described: mate choice and putative genetic incompatibilities. We find that flies from Subtropical Africa harbor previously undescribed genetic diversity and population genetic structure. We also find that *D. melanogaster* has experienced a complex history of gene flow, particularly within Africa. Finally, we find that behavioral isolation and hybrid incompatibilities segregating within the species have a multilayered history, but overall, do not explain patterns of population genetic structure in Subtropical Africa. These results contribute not only to our understanding of the natural and evolutionary history of a powerful model system, but also in our understanding of how reproductive isolation influences global patterns of population genetic structure. We discuss each of these implications in turn.

### Population genetic structure in the ancestral range of *Drosophila melanogaster*

*Drosophila melanogaster* individuals from its purported ancestral range of Subtropical Africa show the highest genetic diversity and the lowest values of Tajima’s D, in line with a demographic history of population expansion and/or strong purifying selection (in agreement with [13,15,16,20,22]). While these flies, originating largely from Zimbabwe, Zambia, Malawi or Namibia, cluster in a genome-wide PCA, our phylogenetic analyses indicate that Subtropical African individuals comprise a non-monophyletic clade that is most distantly related to all other *D. melanogaster,* as would be expected if the mopane forest in countries like Zambia and Zimbabwe are the likely origin of *D. melanogaster* variation [4, 5]. However, we also describe a strong signal of structure among individuals from the ancestral range. Of the eight ancestries identified, four are most prevalent within Subtropical Africa, only one of which is monophyletic in our phylogeny. This monophyletic ancestry group—which we refer to as Harare Distinct (HD) —comprises a distinct cluster in a whole-genome PCA (Figure 1). HD has substantially reduced nucleotide diversity and elevated Tajima’s D, indicative of a population bottleneck or recent introgression (or less likely, balancing selection; [59]). These individuals also have evidence of elevated introgression with OOA relative to Southern Africa. Given that seven of nine individuals in this lineage are derived from the urban center of Harare, HD may be a product of human-assisted migration of OOA individuals into urban centers in Subtropical Africa. This is in agreement with previous findings based on microsatellites [25], as well as previously identified patterns of human-assisted migration in West Africa [26]. While our results partially agree with this hypothesis, we also demonstrate that HD is unlikely to simply be an admixed population between OOA and the Southern African lineage. Although much remains unknown about the HD clade, the presence of previously undescribed diversity within the ancestral range of arguably one of the most well studied model organisms highlights the importance of thorough sampling from ancestral ranges when quantifying diversity and structure.

*Drosophila melanogaster* from other regions within Africa also show strong population genetic structure, with West Africa, East Africa, and Ethiopia harboring three distinct ancestry types, each clustering uniquely in a PCA, and West Africa and Ethiopia being largely monophyletic. Individuals from these areas also have intermediate levels of diversity and Tajima’s D. This supports the hypothesis that these flies likely represent some of the first steps of range expansion from Subtropical Africa (as suggested by [5]). We also find evidence of gene flow between Subtropical Africa and both East and West Africa. In contrast, Ethiopia does not show any introgression with Subtropical Africa and represents a genetically unique lineage from other East Africa flies (in agreement with [16, 18]). Many of the Ethiopian samples herein originate from the high elevation populations from the Ethiopian Highlands, which may be an environment that is not permissive to migration from the lowlands [60–62]. Substantial geographic and/or ecological barriers to migration may thus isolate individuals from the Ethiopian Highlands from the rest of Africa.

In contrast, individuals from outside of Africa show substantially reduced genetic complexity. All OOA flies cluster together in a genome-wide PCA, and largely comprise a single ancestry. As a whole, these samples have reduced diversity and elevated Tajima’s D, which is suggestive of a bottleneck, likely accompanying an out of Africa expansion (as has been found previously [13,20,22,24,63] and reviewed in [64]). We find that all OOA individuals are sister to a subset of East African individuals, specifically those from Kenya, but as a whole, OOA lines do not show evidence of introgression with East Africa. It is therefore possible that OOA lineages are derived from East Africa, and specifically from the ancestors of modern Kenyan flies. Additionally, we find that African lineages have differentially contributed to genetic variation within OOA via post-expansion introgression. We find that all OOA flies have similar levels of introgression with the Southern Africa and HD lineages, while individuals from the SE United States show slightly elevated levels of gene flow with flies from West Africa, relative to other OOA individuals. It has previously been suggested that flies from the SE United States represent a secondary contact zone between individuals of European and African ancestry [16,18,27,45]. Here we show that African lineages have contributed differentially to *D. melanogaster* populations outside of Africa.

Taken together, these broadscale analyses indicate that *D. melanogaster* has substantial population genetic structure within the ancestral range in Subtropical Africa, including the presence of a unique, and potential admixed lineage found in Harare, Zimbabwe. Patterns of gene flow within Africa are complex, as there is at least some degree of gene flow among most genetic lineages, with highland Ethiopian lines remaining relatively distinct and unconnected. Gene flow between African and non-African lineages is also variable, with Subtropical and West Africa exhibiting the highest levels of gene flow with OOA, and West Africa specifically exhibiting a unique pulse of gene flow with flies from the SE United States.

### Introgression is elevated in genes related to human-commensalism

In addition to variation in broadscale patterns of gene flow between distinct genetic lineages of *D. melanogaster,* we also find that patterns of introgression vary across the genome. First, using *f*_dM_, introgression is higher on the *X* chromosome than autosomes for all comparisons (Table S4). This is unusual given that sex chromosomes are often depleted for introgression [65–70]; indeed, within *D. melanogaster* OOA populations often lack African ancestry on the *X* (for example see [45]), although several examples of systems with high levels of gene flow on the *X* also exist [71–74]. Because the *X* chromosome also contains an excess of alleles associated with Z/M female mate choice, our discovery that the *X* chromosome exhibits the highest levels of introgression has implications for the efficacy of female mate choice as a barrier to introgression, a possibility that we explore below.

Genomic windows with the highest levels of introgression contain an excess of genes associated with human commensalism, specifically genes involved in insecticide resistance, metabolism/diet, and immune system function. Human commensalism is a novel, and often adverse environment for organisms. Human-induced habitat change is a primary driver of hybridization in many systems [75, 76]. While the loss of local adaptation due to introgression of domesticated traits is often considered to be a cost of hybridization [77], wild populations without certain domesticated traits may simply be unable to survive in human proximity. The genes we identify as outliers in introgression analyses are often associated with the unique pressures of human commensalism (see [78] for an overview)— insecticide resistance [5,79,80] and diet/metabolism [81–85] are common targets of selection via human commensalism. Increased population densities, and/or novel parasites and diseases associated with human commensalism may also favor increased or novel immunological function [86]. Previous work has established that *D. melanogaster* from non-human associated locales show elevated evolution of some immunity-related genes [5], suggesting that the transition to human commensalism in *D. melanogaster* specifically may involve allelic changes in immunity.

We also find that several genes involved in mate choice and courtship behavior fall within the top 1% of introgressed regions. One potential explanation is that strong female mate preference comes at a fitness cost in more resource-competitive urban environments, potentially by limiting mating potential or delaying reproduction. Thus, it is possible that the cost:benefit ratio of female choice may differ under scenarios of human commensalism, ultimately favoring the loss of female choice in these environments. Under this hypothesis, subsequent introgression of less-choosy alleles back into more urban regions of Subtropical Africa may be favored. Although much more work is needed to further test this hypothesis, previous work has demonstrated the loss of female preference in more urban populations of *D. melanogaster* from Brazzaville, Congo [26]. Similarly, we find that the potentially admixed urban HD clade contains fewer isofemale lines with female mate choice than do the rest of our sample from Subtropical Africa, despite close geographic proximity (Figure 4, Figure S5).

### Female mate choice is polymorphic and likely highly polygenic

Female mate preference in Subtropical African *D. melanogaster* is a classic example of incipient speciation [31–33,35,87–89]. We revisited this classic system to survey flies from Subtropical Africa for the presence and strength of female mate choice, particularly in previously unsampled and remote locales in Zambia, Zimbabwe, and Namibia. We find that female mate choice is relatively common, but also highly polymorphic within and among sampling locales. Female mate choice is also not monophyletically distributed. These results are in line with previous work that found substantial variation in female mate preference for a smaller number of isolines [35], as well as recent work highlighting the variation in male courtship behavior within Subtropical Africa [31– 33,35,87–89]. Three evolutionary scenarios may explain this polymorphism: (1) female mate choice is an ancestral polymorphism which is maintained in Subtropical Africa, (2) strong female mate choice is the ancestral condition, and introgression from outside of this region has eroded its prevalence within Subtropical Africa, or (3) female mate choice is a derived character within Subtropical Africa and has either been eroded by introgression or never fully fixed throughout this region. While we cannot disentangle these scenarios with our current dataset, unraveling these potential scenarios has implications for the stability of reproductive isolation in natural populations, as well as for the potential evolutionary costs and benefits of female choosiness.

One potential caveat to the above female mate choice work is that we used standard Z and M lines for all assays as a way to standardize the female preference tests. However, one possibility is that female mate choice is more common than we estimate, but preference is specific to local males (e.g. not all Z males are equivalent; [87–89]). Several lines of evidence suggest that this may be true for some populations. For example, the population with one of the highest levels of female preference was the population from which the focal male was derived (e.g. ZS). Additionally, studies that used a standard M line and a male of the same isoline as the focal female generally find greater values of female mate choice [35]. Lastly, African males vary in both cuticular hydrocarbon profiles and mating displays, which may be key traits involved in female mate choice [89, 90]. Therefore, while we present the most widespread survey of female mate choice from Subtropical Africa to date, more research will be needed to more precisely characterize the dynamics of female choice in this system.

We leveraged natural variation in the presence of female mate choice within Subtropical Africa to assess genomic differences associated with female choosiness and male attractiveness. Although genomic outliers exist on all major chromosome arms, chromosomes 3L and the *X* are highly enriched for outliers. These results are largely congruent with previous analyses [31, 32], finding that the third chromosome had the largest effect on both female preference and male attractiveness, including a large contribution from 3L. In contrast, [31], found a much smaller contribution of the *X* chromosome in single chromosome replacement experiments, but substantial epistasis between the *X* chromosome and both major autosomes.

Behavioral *PBE* outliers are significantly enriched for genes that are involved in male courtship behavior, olfactory perception, learning, memory, and neurological development. Several individual genes present interesting candidates for future work, including genes involved in mushroom body development and sensory perception. Mushroom body ablation in virgin females results in higher rejection rates [56, 57], and influences male memory and courtship effort [91]. Additionally, while the precise male traits controlling attractiveness are unknown, cuticular hydrocarbons likely play an important role [87,89,92]; and therefore, evolution of sensory perception—particularly sensory perception of smell and taste—may also help regulate female mate preference (e.g. ([93]). Lastly, we find that two genes (*shep* and *eag*), which have been shown directly to affect mating behavior, are highly differentiated in Subtropical African lines with significant female preference. *Shep* is responsible for neuron remodeling during metamorphosis, and whose loss-of-function results in increased rejection of male mates in virgin females [94]. *Eag* regulates potassium permeability, and mutants of this gene have been shown to influence the amount of time males spend courting [95, 96]. Thus, while we present some intriguing candidate genes for future functional analyses, overall our work suggests that the genetic basis of mate preference is likely highly polygenic.

Lastly, while we find that *PBE* behavioral outliers show elevated differentiation between a broader sampling of the Southern African lineage and non-Southern African populations (particularly those from OOA), we also find that these behavioral *PBE* outliers do not show decreased levels of introgression relative to the rest of the genome, nor do lines with strong female preference show a reduced history of introgression. Taken together, these results could arise from a scenario in which female mate preference has experienced polygenic local adaptation, but asymmetric mate preference is insufficient to noticeably dampen gene flow between mating types in Subtropical Africa. In particular, female mate choice may still be maintained if the genetic basis of mate choice is partially redundant (e.g. multiple alleles are sufficient to cause female mate choice, and therefore the loss of any given allele by introgression does not dampen female mate choice). Earlier mapping efforts [31, 32] suggest the existence of genetic redundancy in female mate preference and male attractiveness, though more detailed genetic analyses are needed.

### Putative incompatibilities are highly differentiated, and many are likely recently derived

We lastly studied the distribution of previously identified putative incompatibility loci across our global sampling of *D. melanogaster.* On average, these loci show slightly elevated differentiation between Southern and West Africa. However, while some pairs of loci have a signature of differentiation within Africa, there are substantially more pairs of loci that are highly differentiated between Europe and both West and Southern Africa. Moreover, we find that of the putative incompatibility loci that are highly differentiated, 10-25% of genes within these windows overlap with our behavioral *PBE* outliers (for within Africa or between Europe and both African lineages, respectively). There are two plausible explanations for the high overlap between these two datasets. First, some of the putative incompatibility loci identified by [45] may be involved in assortative mating [58]. Possible candidates for this scenario in our analyses include *eag,* which has been shown to directly influence courtship behavior; *DAAM*, which is involved in mushroom body development; and *rbp6* which influences male aggressive behavior. Second, while *PBE* analyses are an effective approach to detect branch-specific evolution for a focal population [5, 55], by definition, this statistic cannot differentiate branch-specific evolution that is specific to a trait of interest versus subtle population structure and/or branch-specific evolution of non-focal traits. Therefore, if some incompatibility alleles exist at higher frequencies in isolines with strong female choice relative to isolines with no strong female mate choice, they too may be included in our *PBE* outlier analysis (even though these alleles may not directly influence female mate choice). Regardless, our results suggest that putative incompatibility alleles show meager genetic structure across the range of *D. melanogaster*, and knowledge about the function of these alleles can improve our understanding of barriers to gene flow within this model system.

## CONCLUSION

Our work contributes to understanding of the evolutionary history of *D. melanogaster*, including the distribution of genetic diversity and reproductive isolation within this model organism. We combined population genomics and behavioral assays to assess how genetic and phenotypic diversity is partitioned across the globe in *D. melanogaster*, especially within its putative ancestral range in Subtropical Africa. While we find significant population genetic structuring throughout the range, we also find that polymorphic reproductive barriers are largely decoupled from broader patterns of population genetic structure. Furthermore, this work provides insights into the complexity of genetic changes underlying female mate choice, and highlights many genes of interest for future study.

## MATERIALS AND METHODS

### Sampling, Sequencing, and Variant Calling

We created 244 new *D. melanogaster* isolines from an original 339 wild collected females derived from seven novel locations in Zambia, Namibia, and Zimbabwe using a similar approach to previously described efforts ([5]; see Table S1 for sampling locations; see supplemental methods for details). We then extracted DNA and created whole-genome libraries for each of 174 unique isofemale lines plus 32 advanced generation isofemale lines collected in Malawi, Zimbabwe and Botswana (see Table S1 for details). Individually barcoded pools of ∼10 isolines were then sequenced by the University of North Carolina (UNC) School of Medicine, on either a single lane of an Illumina HiSeq 4000 or a Noveseq6000S4XP platform, in both cases generating paired-end 150 bp reads (see supplemental methods and Table S1 for details). These re-sequenced lines were paired with whole genome sequences for an additional 247 isolines via NCBI SRA, including 72 lines from outside of Africa and 175 from within Africa, 30 of which are derived from Subtropical Africa [16, 18] (see Table S1 for details).

We generated a vcf of all 420 *D. melanogaster* plus 12 *D. simulans* and 1 *D. yakuba* genomes using a standard pipeline that followed best practices (see supplemental methods and Table S1 for details). The resultant VCF was filtered so that indels were removed, and only biallelic sites with a minimum quality score of 30, minimum coverage of 5X, minimum genotype quality of 30, a maximum of 50% missing data were kept. We additionally removed 10 individuals with poor quality genomes (e.g. less than 5X average coverage).

### Lineage relationships: population structure, PCA, and phylogenetic reconstruction

To better understand the relationships among a global sampling of *D. melanogaster*, we constructed maximum likelihood (ML) phylogenies for the autosomes and *X* chromosome separately using *iqtree* version 1.6.12 [97–99]. We generated ML trees for non-overlapping 100KB windows using the model-finder and ultra-fast bootstrap approach with 1,000 bootstraps. We then used the resulting ML trees as input for *ASTRAL v5.1.1* [100] in generating a consensus tree for the autosomes and *X* chromosome independently (Figure S2).

To characterize fine scale population genetic structure we performed *K*-means clustering analysis and PCA using *PCAngsd* [50] for the *D. melanogaster* samples. PCAngsd uses genotype likelihoods to first perform a genome-wide PCA, then assess population structure with the number of ancestry types (*K*) defined as the number of significant PCs + 1. We only included sites with <10% missing data, a minimum mapping quality of 30, and a minimum base quality of 20. This generated a dataset of 461,147 sites for 420 individuals.

Given our phylogenetic, PC, and *K*-means clustering analysis, we identified six genetic lineages which correspond to five geographic regions (all Out of Africa lines (OOA); all East African lines excluding lines from Ethiopia (East), all West African lines, all Ethiopian lines, nine lines from Subtropical Africa which form a unique ancestry group that we refer to as Hahare-Distinct (HD), and all other Subtropical African lines (Southern)). We calculated all pairwise measures of divergence and differentiation between these lineages (e.g. *F*_ST_ and *D*_xy_; defined in Figure 1) and nucleotide diversity within each lineage (e.g. *π*) in 1KB windows across the genome using *pixy* [101] with default filtering expressions (i.e. DP>=10,GQ>=20,RGQ>=20). For this analysis we used an all-sites VCF to include invariant sites, and performed the same filtering above, but retained sites with a maximum of 2 alleles, so as to include invariant and biallelic sites. We also calculated Tajima’s D using *VCFTools* [102], and estimated the Site-Frequency Spectra (SFS) using *SweeD* [103] for each genetic lineage.

### Introgression Analyses

To estimate broad patterns of gene flow between genetic lineages of *D. melanogaster,* we first calculated Patterson’s *D* and *f*_G_ (a Patterson’s *D* derivative which more accurately estimates the proportion of the genome experiencing introgression [104]) using *Dsuite* [54] with *D. simulans* and *D. yakuba* as outgroups for all possible trios, given the following phylogeny: (((OOA, ((East, West),Ethiopia)),HD), Southern). Significance of Patterson’s *D* was determined using a standard block jackknife procedure [54]. In specific cases (outlined below) we also quantified differences in the extent of introgression between genetic lineages that exhibited significant Patterson’s *D*, using *f*_dM_ in non-overlapping 20 SNP windows for focal trios (outlined below). *f*_dM_ is a Patterson’s *D* derivative that is more appropriate for windowed analysis and provides a more accurate estimate of the proportion of the genome that has experienced introgression than Patterson’s *D* [54]. Finally, we bolstered our introgression analyses by assessing heterogeneity in the genome in the relationships between potentially introgressing groups by calculating tree topology weights using *twisst* [105]. For each focal trio (outlined below), we calculated the topology weight at each non-overlapping 100KB window for trees comprising four groups: *D. simulans* as an outgroup, the two potentially introgressing groups (outlined below) and the remaining African flies as a single group (AFR). While *f*_dM_ calculates the proportion of shared derived variants between non-sister lineages, *twisst* assesses the proportion of topologies that fit particular phylogenetic relationships. These analyses thus provide complimentary, but uniquely informative quantifications of introgression.

Our goals were threefold: (1) assess if female mate choice in Subtropical Africa is associated with reduced gene flow with other African lineages, (2) quantify if urbanization in Subtropical Africa is associated with increased levels of gene flow with OOA, and (3) identify the source(s) of African ancestry in the SE United States. To address each of the specific questions listed above, we used specific trios.

#### Gene flow between Southern Africa and other African lineages

we first assessed patterns of gene flow between all trios involving HD or Southern Africa as P3, and Ethiopia, West, and/or East Africa as P1 and P2 using Patterson’s *D*. For trios that had significant Patterson’s *D*, we then calculated *f*_dM_ in 20 SNP windows across the genome to evaluate the extent of introgression. To determine whether the extent of introgression varied between trios, we used an ANOVA with windowed *f*_dM_ as the response variable, and P3 (HD or Southern Africa), P2 (East or West), and their interaction as independent variables with a Type III Sum of Squares using the *car* package in R [106]. In all cases, Ethiopia was used as P1, as there were no scenarios in which Ethiopia demonstrated significant introgression with either HD or Southern Africa (Table S3; Figure 3).

#### Urbanization and introgression with OOA

we calculated Patterson’s *D* for all trios in which Southern Africa (composed largely of rural individuals) or HD (composed largely of urban individuals) was P3, and OOA and either Ethiopia, East, or West Africa was P1 or P2. Then, to determine if either HD or Southern Africa has experienced more introgression with OOA, we calculated *f*_dM_ in 20 SNP windows across the genome for both comparisons with Ethiopia as P1, OOA as P2, and either Southern Africa or HD as P3. We assessed differences in windowed *f*_dM_ using an ANOVA with Type III SS using the *car* package in R [106], with *f*_dM_ as the independent variable and P3 (HD or Southern Africa) as the independent variable.

#### Source(s) of African ancestry in the SE United States

we performed two sets of analyses. Again, we calculated Patterson’s *D* with either lines from the SE United States or all other OOA lines as P3, and Ethiopia, East and/or West Africa as P1 and P2. We calculated Patterson’s *D* with HD or Southern Africa as P3 and either lines from the SE United States or all other OOA as P2, and each of Ethiopia, East and West Africa as P1. As flies from the SE United States are hypothesized to represent a contact zone between OOA and an unidentified African lineage, we expect that flies from the SE United States should have increased levels of introgression with the source African lineage(s), relative to the rest of OOA flies. Therefore, we again calculated *f*_dM_ in 20 SNP windows across the genome for trios with significant Patterson’s *D*, and asked which (if any) African lineages had higher *f*_dM_ with the SE-United States than with OOA using ANOVAs with Type III SS using the *car* package in R [106] with windowed *f*_dM_ as the response variable and each OOA lineage (SE-United States or all other OOA lines) as the independent variable.

Last, to assess heterogeneity in introgression across the genome, we leveraged the windowed *f*_dM_ analyses described above and first asked if chromosomes varied in the extent of introgression using an ANOVA with Type III SS in the *car* package in R [106]. For each focal trio we performed these analyses with *f*_dM_ as the response variable and chromosome arm as the independent variable. We then used pairwise *t*-tests to assess significant differences between chromosome arms. To determine what genomic regions were specifically more likely to introgress in each of our focal comparisons, we also defined introgression outliers as windows with the top 1% of *f*_dM_ values.

### Assessing mate preference in Subtropical African samples

To assess the prevalence and strength of female mate choice in Subtropical Africa, we performed a series of replicated choice experiments for 47 focal isofemale lines from Subtropical Africa. Briefly, each choice experiment was conducted by presenting a 7-10 day old virgin female with a standard Z and M male line (ZS2 and RAL371, respectively). Vials were watched continually for up to three hours, and once mating began the unmated male was removed from the vial by aspiration. We completed an average of 53 trials (range: 12-100) per focal isofemale line, with an average of 37 trials resulting in a successful mating per isofemale (range: 8-71; 1,724/2,507 total trials were successful). See the supplemental methods for full experimental details.

We assessed deviations from random mating for each isofemale line using a Fisher’s exact test, then assessed population-level differences in the proportion of females exhibiting preference and the strength of preference using a logistic regression or a generalized linear mixed effect regression with poisson distribution, respectively. In both models, population was the independent variable. For the model assessing the strength of preference, we also included the isoline, replicate trial date, and total matings as random effects. For both of these analyses we used the *lme4* package [107] and assessed significance of fixed effects using an Type III Wald’s X^2^ test using the *Anova* function in the *car* package in R [106].

To test for genetic differences associated with female mate preference, we first calculated the Population Branch Excess (*PBE*) statistic [55] in 1KB windows across the genome with Subtropical African lines with significant female mate choice as the focal population, and Subtropical African lines that do not have strong female mate choice and RAL lines as the non-focal populations. *PBE* is a derivative of the population branch statistic (i.e. *PBS*; [108]), but quantifies branch-specific evolution for a focal population relative to two non-focal populations [55]. We then defined behavioral *PBE* outliers as the top 1% of *PBE* values (368 windows). Using this list of outliers, we did a Gene Ontology (GO) Enrichment Analyses with a Fisher’s Exact test with an FDR threshold of 0.05 using *PANTHER* 16.0 [109] to assess if behavioral *PBE* outliers were enriched for biological GO terms relating to behavior, sensory perception, memory, learning or neurological development, each of which may have specific function in female choosiness or male attractiveness [96].

Lastly, we aimed to determine whether female mate choice in Subtropical Africa is associated with global patterns of differentiation or introgression in *D. melanogaster* to assess whether female mate choice serves as a significant barrier to gene flow in nature. To do this, we performed three subsequent analyses. First, we calculated whether behavioral *PBE* outliers had elevated differentiation relative to the rest of the genome between the Southern Africa lineage and every other genetic lineage. We used the aforementioned estimates of *F*_ST_ calculated in 1KB windows between all Southern Africa lines (regardless of phenotype) and every other genetic lineage. We then performed an ANOVA with Type III SS to assess whether *F*_ST_ differed between locus type (genome-wide versus behavioral *PBE* outlier), between comparisons (each genetic lineage), and their interaction using the *car* package in R [106]. We used the *emmeans* package to estimate significance of specific contrasts [110]. Second, to determine whether loci potentially involved in female mate choice were associated with differential patterns of introgression, we also ascertained whether windowed *f_dM_* between Southern Africa and each of West Africa and Europe differed between behavioral *PBE* outlier loci and the rest of the genome, using the *f_dM_* datasets described above. To do this, we performed a *t*-test with windowed *f_dM_* as the dependent variable, and locus type (behavioral *PBE* outlier vs genome-wide) as the independent variable. Third, we asked whether isolines that had significant female mate choice also showed an overall lower history of introgression than isolines with no strong female mate choice. To do this, we recalculated Patterson’s *D* and *f_G_* for trios involving OOA, West Africa, East Africa, and/or Ethiopia as P1 and P2, and isolines of each behavioral type as P3. We restricted these analyses to include only Southern Africa flies for which we had known behavioral phenotypes, and excluded phenotyped flies from the HD lineage, as these genetic lineages exhibit substantial population structure (Figure 1; Figure S5). We then performed a *t*-test with *f_G_* as the dependent variable and the identity of P3 (i.e. lines that showed behavioral preference versus those that did not) as the dependent variable.

### Determining the global distribution of previously identified incompatibilities

We characterized patterns of differentiation for loci that have previously been implicated in two studies of genetic incompatibilities with *D. melanogaster* [45, 46]. First, [46] used a global panel of *D. melanogaster* inbred lines to create synthetically admixed populations from a series of round-robin matings followed by continual inbreeding. This design enabled the identification of pairs of alleles that appear less frequently than expected under random mating and Mendelian segregation in their final recombinant inbred line population (i.e. Genotype Ratio Distortion). Using, a similar premise, but in a naturally admixed population, Pool [45] used patterns of linkage disequilibrium in the SE United States to assess pairs of alleles that occur together less frequently than expected based on their allele frequencies (i.e. Ancestry Disequilibrium). [45] also determined that many of these loci were highly differentiated between Africa and Europe, using populations from West Africa and France, respectively.

Elevated differentiation of these putative incompatibility alleles between West Africa and France may stem from multiple evolutionary scenarios, and differentiating these scenarios can help elucidate the geographic distribution and potential origins of putative incompatibilities within *D. melanogaster*. Here, we aim to differentiate two potential scenarios: First, putative incompatibilities between Europe and Africa may have arisen with or after the Out of Africa expansion and thus may represent more recently derived incompatibilities (i.e. within the last 10-23kya; [5,20,22]). Under this scenario, we predict that differentiation at putative incompatibility loci should be low between genetic lineages in Africa, but high between Europe and all African populations. Second, it is also plausible that putative incompatibilities between Europe and West Africa are very old and also exist between West Africa and Southern Africa, with shared ancestry or subsequent introgression explaining allele sharing between Europe and Southern Africa. Under this scenario, we expect that differentiation should be high between West Africa and both Europe and Southern Africa, but relatively low between Europe and Southern Africa.

To differentiate these scenarios, we used the *F*_ST_ windows from above for all pairwise comparisons of Southern Africa, West Africa, and Europe. We first ask if putative incompatibility loci have elevated divergence relative to the whole genome for any comparison using an ANOVA with Type III SS using the *car* library in *R* [106]. Specifically, *F*_ST_ was the dependent variable, and locus type (genome-wide, loci identified by [46], or loci identified by [45]), population pair (Southern Africa-West Africa, Europe-Southern Africa, or West Africa-Europe), and their interaction were the independent variables. We then used the *emmeans* package in R [110] to determine significance for specific contrasts. Second, we assessed the history and distribution of individual pairs of loci and differentiate the two evolutionary scenarios outlined above. For these analyses, we identified putative incompatibility pairs in which both interacting loci have the predicted patterns of differentiation. We define highly differentiated loci as those with *F_ST_* values within the top 2.5% of *F_ST_* for that population pair.

## Supporting information

Supplemental Methods_TableCaptions_+Figures

## ACKNOWLEDGEMENTS

We graciously thank the past and present members of the Matute lab, as well as Kieran Samuk for thoughtful discussions on this project. This work was funded by an NIH R01 to DRM (R01GM121750) and an NIH R35 (R35GM124701) to BSC.

## DATA AVAILABILITY

All new raw sequence data will be made available on the NCBI SRA upon acceptance of this manuscript. Current SRA codes are noted in Table S1. Raw behavioral data will be uploaded to Dryad upon acceptance of this manuscript.

