## Supplemental Methods_TableCaptions_+Figures for "Patterns of and processes shaping population structure and introgression among recently differentiated *Drosophila melanogaster* populations"

#### *Sampling*

We collected *D. melanogaster* from seven locations in Zambia, Namibia, and Zimbabwe using a similar approach to previously described efforts ([5]; see Table S1 for sampling locations). Our approach differed from other samplings in that we used multiple potential substrates. Each trap consisted of buckets adjacent to each other and separated by about 50cms. The buckets were filled with mashed bananas (purchased locally), Marula (*Sclerocarya birrea*) or muzinzila fruits (*Berchemia discolor).* In all cases, we removed the husk of the fruit, added yeast (Red Star Active Dry Yeast - 16 oz. #201265), and allowed them to ferment for about 24 hours. These traps were put underneath trees. We collected all flies in the bucket using a sweeping net (BioQuip; Rancho Domingo, CA) after 24, 48, and 72 hours. We then netted and aspirated flies with a pooter (1135A Aspirator–BioQuip; Rancho Domingo, CA)) and anesthetized within 20 minutes of collection using FlyNap (triethylamine, Carolina Biological Supply Co.). Females and males were separated. Males and individuals from other species were placed in ethanol; *D. melanogaster* females were placed in 30mL plastic vials with cornmeal food and allowed to oviposit. Of 339 collected females, we were able to establish 244 isofemale lines (i.e., shelf-stable lines derived from a single matriarchal lineage).

#### *DNA extraction and sequencing*

We extracted DNA from 20 individuals from each of the 174 unique isofemale lines using a Gentra Puregene Tissue Kit (Qiagen, Valencia, CA, USA) following the recommended tissue protocol with volumes of reagents as suggested for processing 5 - 10 mg of tissue (see Table S1 for collection details). To prepare the genomic DNA libraries we used KAPA HyperPrep kits (Roche Sequencing, Pleasanton, CA) with a target fragment size of 300-500 bp at the University of North Carolina (UNC) School of Medicine's high-throughput sequencing facility. Next, we pooled individually barcoded libraries into groups of ~10 individual libraries and each pool was sequenced on either a single lane of an Illumina HiSeq 4000 or a Noveseq6000S4XP platform, in both cases generating paired-end 150 bp reads. This sequencing strategy yielded between 2.6-23.1 billion bp of raw sequence data for each individual (See Table S1 for coverage information). Additionally, we sequenced 32 lines that were advanced generation isofemale lines collected in Malawi, Zimbabwe and Botswana (outlined in Table S1).

*Public data*

We obtained whole genome sequences for an additional 247 isolines via NCBI SRA. Of these publicly available genomes, 72 were of flies sampled outside of Africa and 175 were of flies collected from within Africa, 30 of which resided in the preported ancestral range in Subtropical Africa [16,18] (see Table S1 for details).

*Variant calling*

We aligned 420 genomes of *D. melanogaster* to the *D. melanogaster* v6.32 reference genome [111] using *bwa mem* function [112]. We then used *Picard Tools* (http://broadinstitute.github.io/picard/) to clean, sort and dedupe individual files before individually genotyping them in *GATK4* [113] with the *HaplotypeCaller* function. All samples were then jointly genotyped in *GATK4* using the *GenotypeGVCfs* function, following *GATK* best practices [113]. The resultant VCF was filtered so that indels were removed, and only biallelic sites with a minimum quality score of 30, minimum coverage of 5X, minimum genotype quality of 30, a maximum of 50% missing data were kept. We additionally removed 10 individuals with poor quality genomes (e.g. less than 5X average coverage). For analyses requiring an outgroup (such as our phylogenetic reconstruction, outlined below), we also included twelve *D. simulans* and one *D. yakuba* genomes (see Table S1 for SRA accession numbers).These sequences were processed as above, and VCF files were merged using *bcftools merge* function [114]. We used this VCF file to perform the population genomic analyses, outlined below.

*Assessing mate preference in Subtropical African samples*

We next tested if patterns of diversity, differentiation, and population structure correspond to Z and M mating types within *D. melanogaster* (as described in [31,33]). While Z mating behavior seems largely restricted to Subtropical Africa, previous work has identified that female mate choice is likely variable in the ancestral range of *D. melanogaster [35]*—however, the frequency of Z behavior within the ancestral range, as well as it's phylogenetic distribution among Subtropical African flies, is largely unknown. To assess the prevalence and strength of female mate choice in Subtropical Africa, we performed a series of replicated choice experiments for 47 focal isofemale lines. These lines originate from four sites in Zambia (a total of 17 lines), three sites in Zimbabwe (a total of 21 lines), one population from Malawi (two lines), one population from Botswana (one line), and one population from Namibia (six lines). Each focal isofemale line was tested in a choice experiment where 7-10 day old virgin females were presented with a standard Z and M male in a vial containing cornmeal/Karo/agar medium. We used the lines ZS2 and RAL371 as representative Z and M lines, respectively. We note that while RAL lines have mixed African-European ancestry, RAL371 is 81.2% European ancestry, slightly above the average 80.2% for the population as a whole [45]. Vials were watched continually for up to three hours, and once mating began the unmated male was removed from the vial by aspiration.

As Z and M males are morphologically indistinguishable, males were transferred to vials containing cornmeal/Karo/agar medium that had been dyed with either red or blue food safe dye approximately one hour prior to the mate choice experiment, and were easily distinguished based on abdominal coloration. Across replicates, we switched which males were fed what color to control for any bias abdominal coloration might have on female preference, and we found no effect of food color on female preference (based on an ANOVA with type III SS, with proportion of Z males chosen as the response variable, and isoline and color of Z male as fixed effects: effect of color of Z male: $F$=0.088, $df$=1, $p$=0.77; effect of isoline: $F$=1.57, $df$=47, $p$=0.039). We tested for female mate preference using a Fisher's exact test to determine if the observed ratio of Z:M successful matings significantly deviated from the expected under random mating (e.g. 1:1) for each isofemale line (we note that the results are qualitatively the same when mate preference is assessed using a binomial test).

### SUPPLEMENTAL TABLES LEGENDS (see attached excel):

**Table S1: Sample information for all individuals in this study.** For each accession, the unique ID (sample), species (each are species names of the genus *Drosophila*), NCBI SRA code (new samples will be given a unique code upon data upload), the sequencing platform from which the data came, the genetic lineage which the sample belongs to, collection location information, and the average depth of coverage are given

**Table S2:** $F_{ST}$ (blue; lupper triangle), $D_{xy}$ (green; lower triangle), and π (diagonal) for all genetic lineages of *D. melanogaster* and *D. simulans*.

**Table S3:** *D* statistics of focal comparisons. Population codes: ET= Ethiopia, East= East Africa, West= West Africa, SE-US= SE United States, SOU= Southern Africa, HD= Harare Distinct, OOA= Out of Africa (excluding SE North America).

**Table S4:** Mean $f_{dM}$ values for each chromosome arm by each focal comparison. Significant differences between chromosome arms for each comparison are indicated by superscript letters.

**Table S5: Introgression Outliers.** Top 1% of $f_{dM}$ outliers for each of three comparisons: West Africa and Southern Africa (Southern-West), HD and OOA (OOA-HD), and West Africa and the SE United States (West-SE-US). Genes within the outlier window and those proximal (as well as their proximity in KB) are given.

**Table S6: Summary of behavioral survey.** Average proportion of Z males chosen across trials for each isoline phenotyped in our behavioral survey, as well as the total trials set up, the total matings recorded, and the proportion of successful trials (mating proportion). We then defined each line as a behavioral type (no preference= NP; preference= P) based on both a Fisher's Exact and binomial test (these analyses agreed completely on type definition with a significance cutoff of $p<0.05$).

**Table S7: Behavioral *PBE* outliers.** Top 1% of *PBE* outliers with Southern African lines that show significant mate preference as the focal population, and Southern African lines with no strong mate preference and RAL lines as the non-focal populations. Names of genes directly within the outlier window, as well as names of genes in close proximity are listed for each window, along with their proximity in KB up/downstream.

**Table S8: Behavioral *PBE* GO analyses.** GO analyses of the genes in or proximal to *PBE* outliers. Performed using *PANTHER* v.16.0.

**Table S9: Putative incompatibilities with $F_{st}$ outliers.** A list of putative incompatibilities identified by [45] and [46] in which both interacting loci show signatures of older or more recent derivation (denoted in the Popgen signature). Window positions were translated between V.5 and V.632 of the *D. melanogaster* reference genome, as different reference genomes were used between [45] and the current study. As some windows identified by pool contain multiple $F_{ST}$ outliers, the number of outliers contained for each locus window, plus the genes that fall within that outlier are listed.

### SUPPLEMENTAL FIGURES:

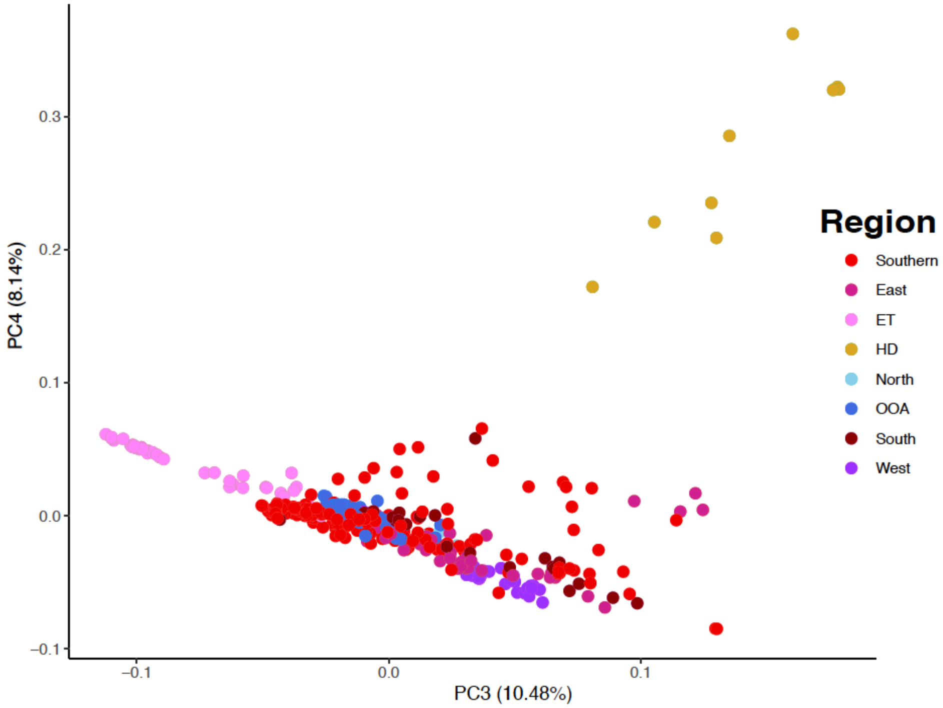


**Figure S1: PCs 3 and 4 of a genome-wide PCA.** *Points are colored based on the genetic lineage. Percent of variance explained by each PC is indicated in brackets.*

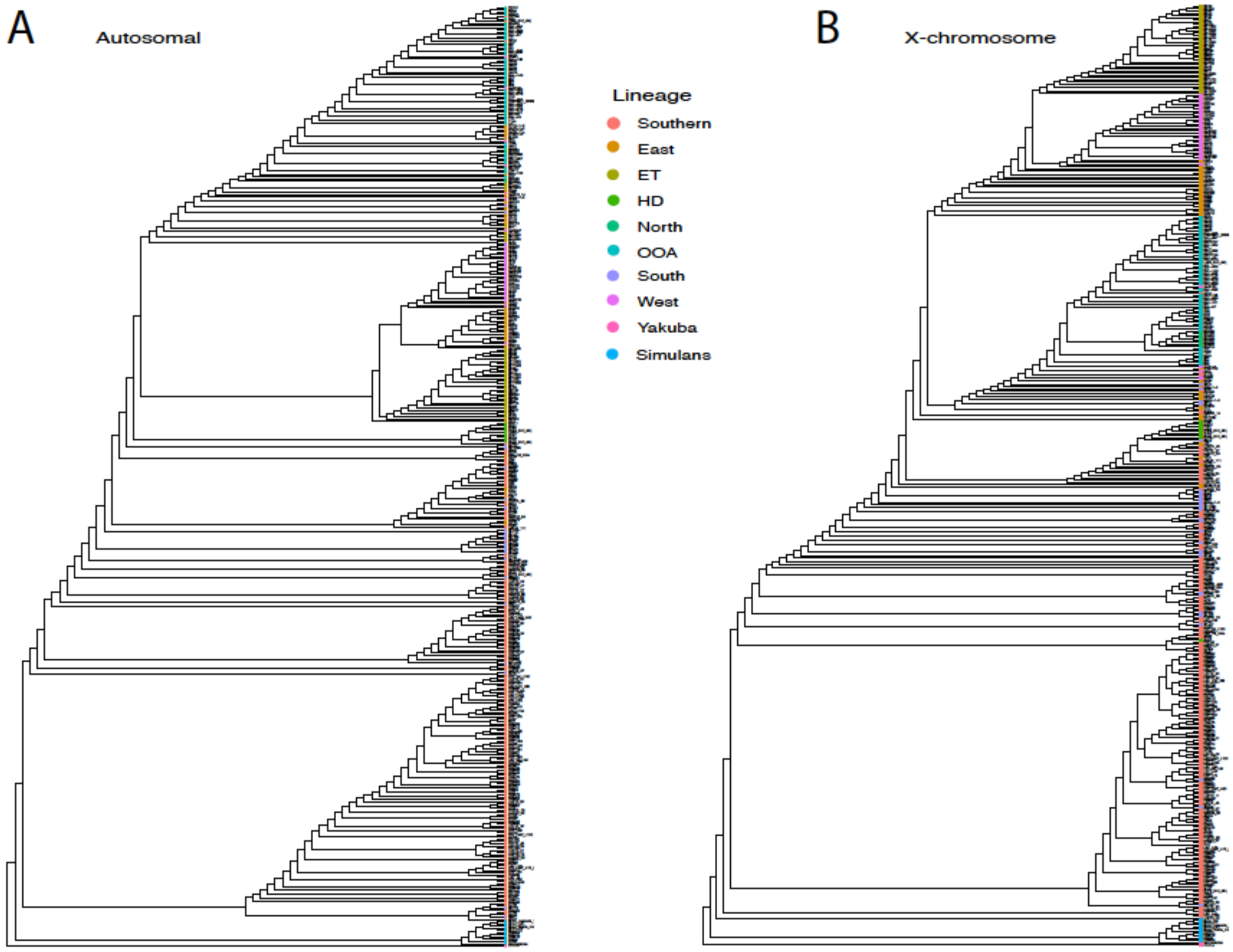


**Figure S2: Full ML phylogeny of all samples** based on (A) autosomes and (B) the *X* chromosome only. Tip colors correspond with the genetic lineages defined herein, plus *D. simulans* and *D. yakuba* as outgroups.

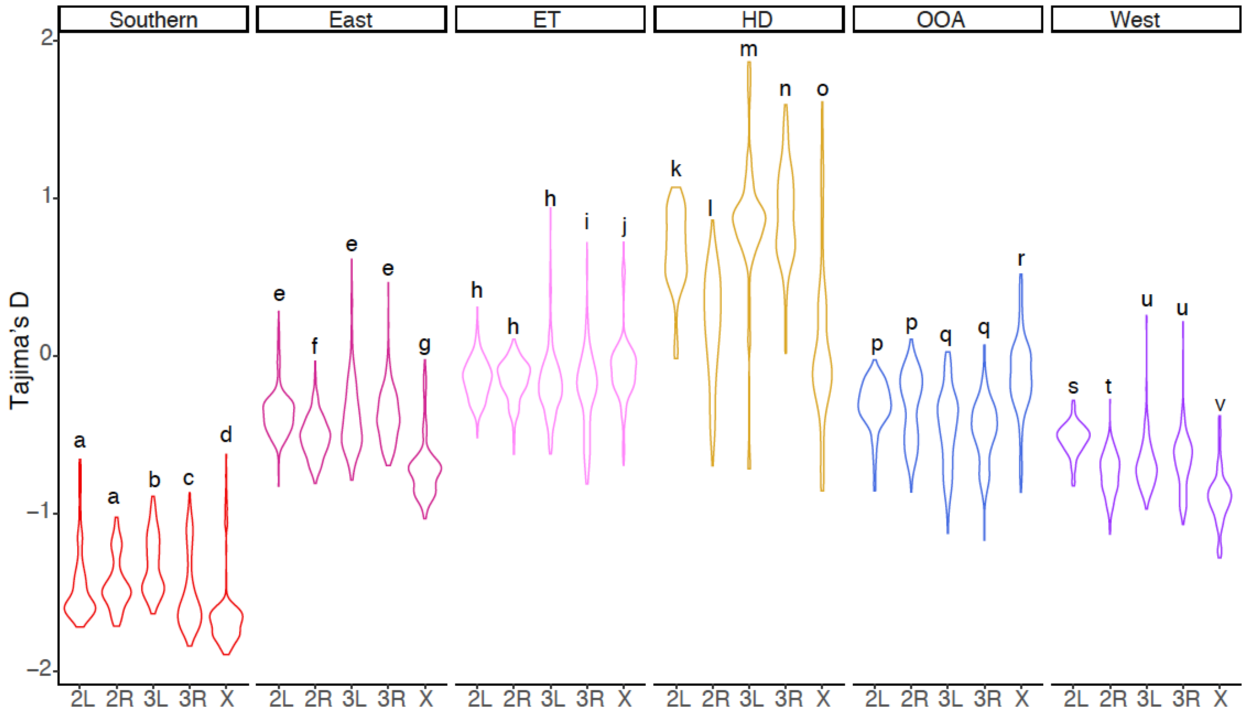


**Figure S3: Tajima's D by chromosome for each genetic lineage.** Letters denote significantly different chromosome arms per lineage.

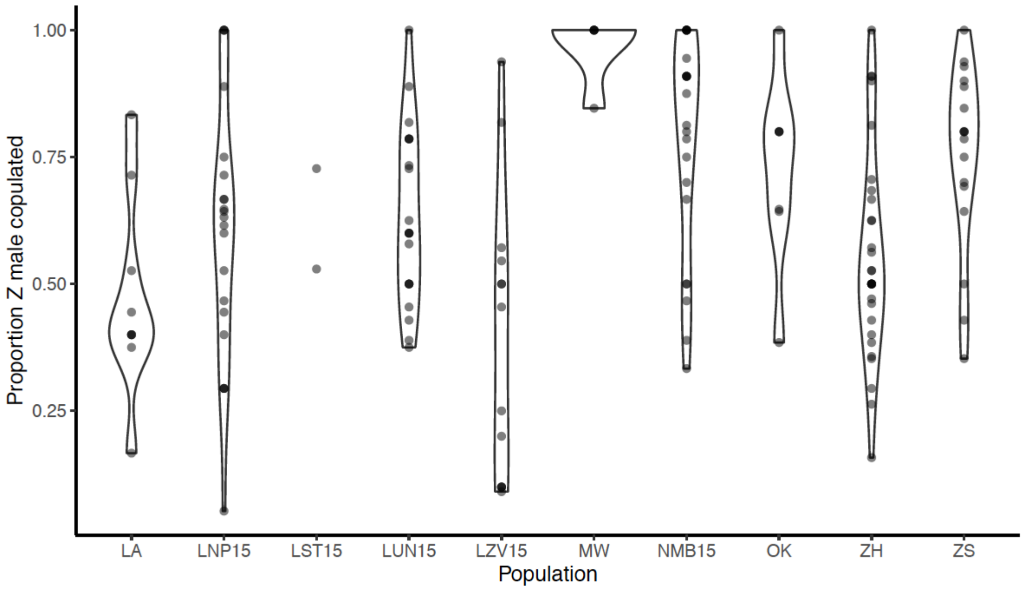


**Figure S4: Strength of female mate preference differs by population.** Proportion of Z males copulated in a choice trial between Z and M males and female isolines from 10 sampling locales in Subtropical Africa. Points represent individual trials for individual isolines.

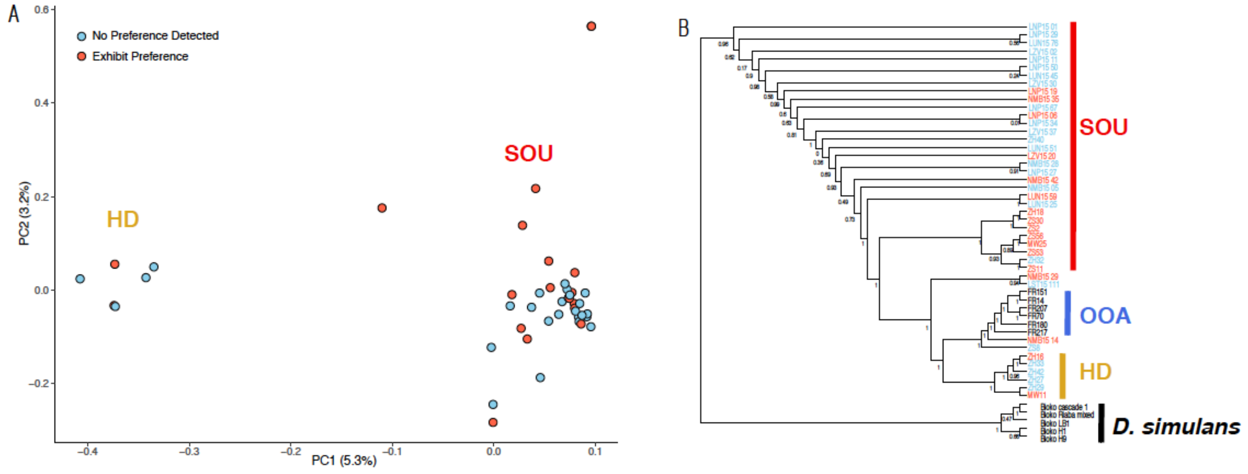


**Figure S5: No genetic structure between lines with or without strong mate preference**; *denoted in red and blue respectively. (A) PCA of all Southern African lines that were phenotype for mate preference. Substantial genetic structure between HD and Southern (SOU) clades, but no structuring between lines that do/do not show significant female mate preference. Moreover, there is no genetic structure between behavioral types across the first 8 PCs. Numbers in brackets denote the percent of variance explained by PC1 and PC2, respectively. (B) ML consensus phylogeny built from non-overlapping regions of 100KB using the rapid-hill climbing algorithm in RAxML, with 50 bootstraps per tree. Consensus phylogeny was built using these trees as input to ASTRAL. Each genetic clade is outlined with a thick vertical bar, with colors replicating the represented clades in Figure 1. We also include six lines from France (denoted by the blue OOA section) for context. We note that only 17 of 19 individuals with strong preference and have sequenced genomes are included in the phylogeny as 2 genomes did not pass quality thresholds for our phylogenetic analyses (LA66 and LA69).*

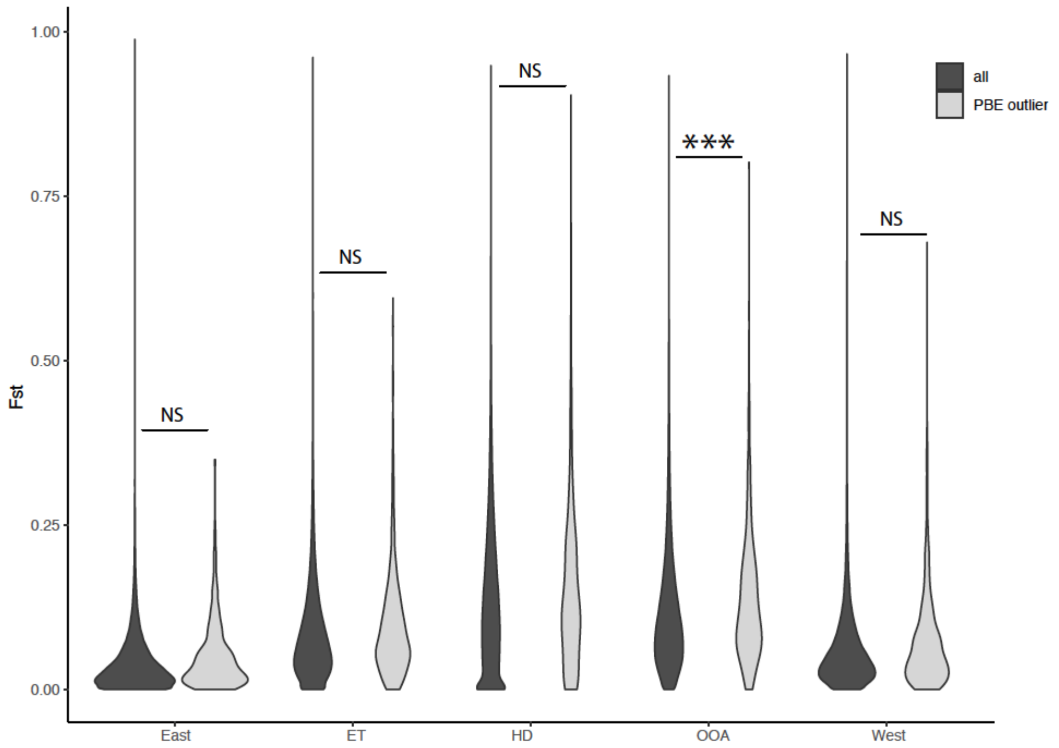


**Figure S6: Behavioral outliers are highly differentiated between Southern Africa and other genetic lineages.** *PBE outliers for behavior show increased $F_{st}$ relative to the rest of the genome for all comparisons (genome-wide versus outlier: F=6.43, df=1, p=0.011). However, when contrasting $F_{ST}$ between Southern Africa and specific genetic lineages, differences between $F_{ST}$ for genome-wide versus PBE behavior outliers is only significant for Southern Africa versus OOA.*

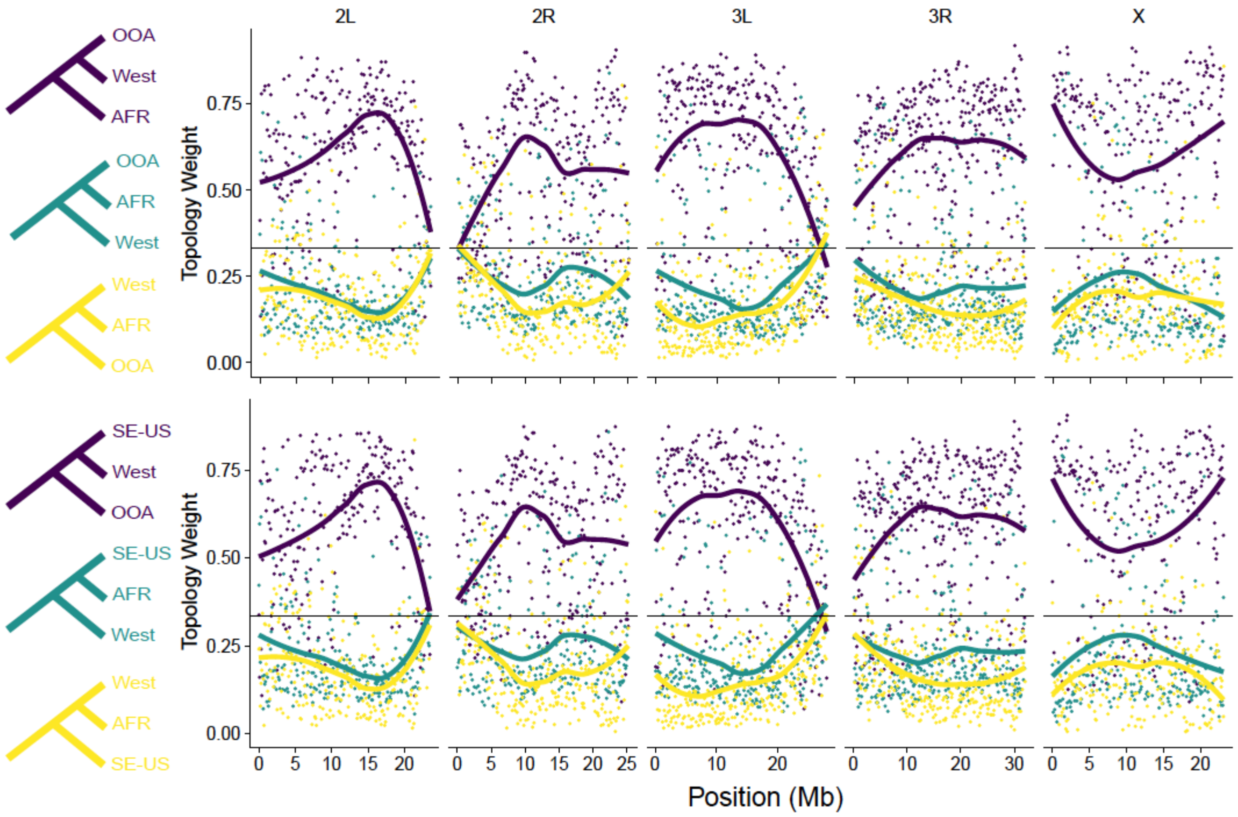


**Figure S7: Weighted topologies between West Africa and OOA.** *Highly similar landscapes of weighted topologies between West Africa and each of all OOA lines (top) and just those from the SE-United States (SE-US; bottom) suggest a largely shared landscape of introgression.*

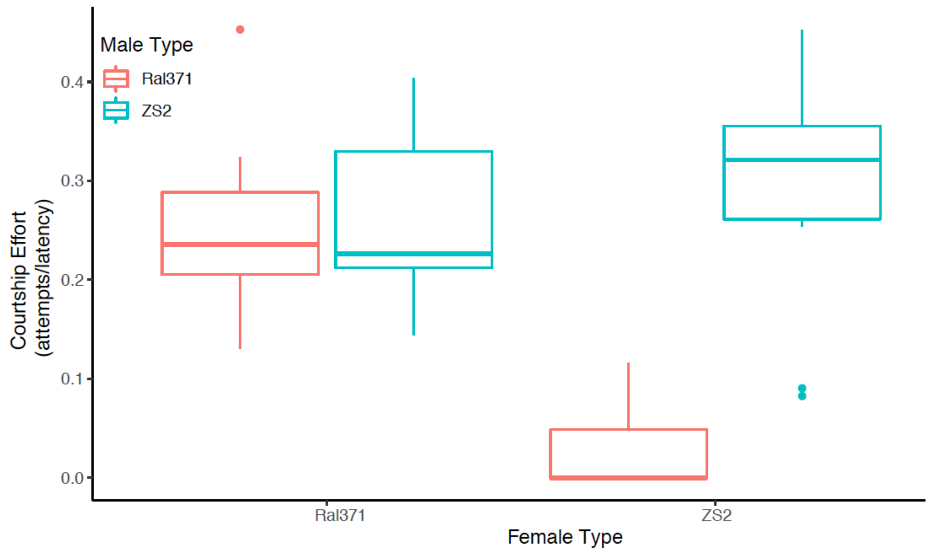


**Figure S8: Courtship effort differs between males of different genotypes.** Courtship effort (defined as the number of courtship attempts divided by latency) per behavioral type of male (M=Ral371, Z=ZS2) for each female in a two-way choice experiment. Z males attempt more courtings, particularly in the context of Z females.

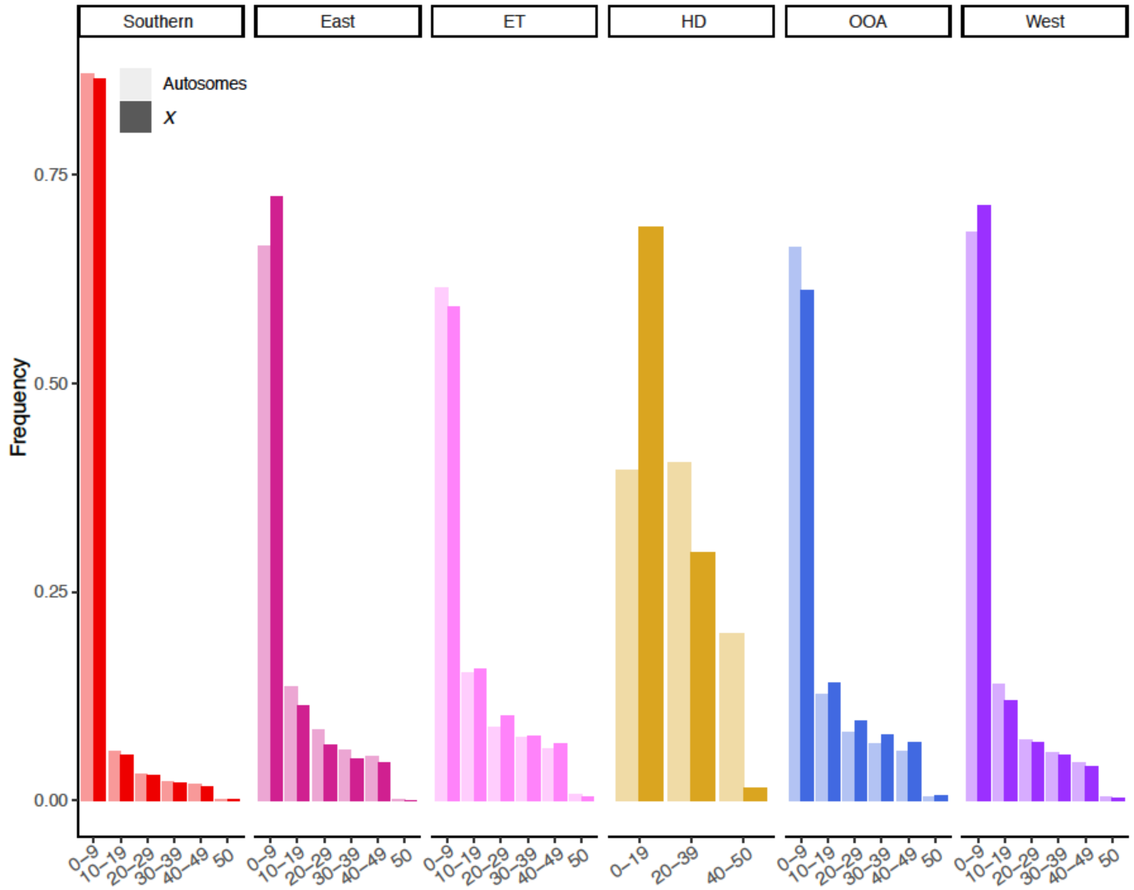


**Figure S9: Folded Site-Frequency Spectrum for each genetic lineage.** Greyed bars are the average across 2L, 2R, 3L and 3R. Note difference in x axis for the HD clade given limited sampling relative to the other lineages.
